# A short linear motif, conserved from yeast to human, binds to members of the Spa2/GIT1 family of cortical scaffold proteins

**DOI:** 10.1101/2025.06.12.659263

**Authors:** Lara Bareis, Annika Siewert, Benjamin Grupp, Tim Bergner, Clarissa Read, Steffi Timmermann, Nicole Schmid, Nils Johnsson

## Abstract

Tip growth is closely tied to fungal pathogenicity. Spa2, a multi-domain protein and member of the polarisome, orchestrates tip growth in yeast and other fungi. We identified a conserved short linear motif in the RabGAPs Msb3 and Msb4, and the MAP kinase kinase Ste7 and Mkk1, which mediates their interaction with Spa2. AlphaFold predictions suggest that these initially unstructured motifs adopt an alpha-helical conformation upon binding to the hydrophobic cleft of Spa2’s N-terminal domain. Altering the predicted key contact residues in either Spa2 or the motif reduces complex stability. Such mutations also cause mis-localization of Msb3, Msb4, and Ste7 within the cell. Deleting the motif in Msb3 or Msb4 abolishes tip-directed growth of the yeast bud. Protein assemblies that spatially confine secretion to specific membrane regions are a common feature of eukaryotic cells. Accordingly, Spa2-motif complexes were predicted in orthologs and paralogs across selected Opisthokonta, including pathogenic fungi and humans. A search for functional motifs in conformationally flexible regions of all yeast proteins identified Dse3 as a novel Spa2-binding partner.

## Introduction

Budding yeast, like other walled fungi, shapes its growth by directing secretory vesicles to specific plasma membrane regions during the cell cycle. The polarisome, a multi-protein assembly, orchestrates this process by selecting target sites for vesicle fusion (Sheu et al., 1998; Sheu et al., 2000; Dünkler et al., 2021). Its primary scaffold protein, Spa2, interacts with two Rab-GAPs, Msb3 and Msb4, which convert the Rab-GTPase Sec4 from its active GTP-bound state to its inactive GDP-bound state (Snyder, 1989; Gao et al., 2003; Tcheperegine et al., 2005). GTP-bound Sec4 recruits the motor protein Myo2 and the exocyst component Sec15 to post-Golgi vesicles (Guo et al., 1999; Jin et al., 2011). The vesicle then moves on actin cables to the sites of the bud membrane where fusion occurs. Deletion of both Msb3 and Msb4 impairs vesicle fusion with the plasma membrane, leading to vesicle accumulation in the bud (Tcheperegine et al., 2005). The GAP activity not only recycles Msb3/4 but also catalyzes a critical, yet poorly understood, step in vesicle targeting or fusion (Gingras et al., 2022). Thus, a key function of the polarisome during tip growth is to localize Msb3/4 to prospective vesicle fusion sites. The interaction between Spa2 and the GAPs occurs via the N-terminal Spa2 homology domain 1 (SHD1) and potentially unstructured N-terminal regions of Msb3/4 (Tcheperegine et al., 2005).

Intrinsically disordered regions, often less conserved evolutionarily, may contain short linear motifs that regulate protein localization, stability, or interactions (Holehouse and Kragelund, 2024; Mughal and Caetano-Anollés, 2025). These motifs are typically recognized by folded domains of their binding partners. Such interactions are often weak or transient, making them challenging to detect. As a consequence, many motifs likely remain undiscovered within the disordered regions of an organism’s proteome (Tompa et al., 2014). However, identifying a motif’s sequence can reveal its presence in other proteins across species, enabling predictions of binding partners, localization, and even cellular roles (Neduva et al., 2005).

Here, we identify a short linear motif (SLiM) in the disordered regions of Msb3, Msb4, the MAP kinase kinases Ste7 and Mkk1 that binds the SHD1 domain of Spa2. AlphaFold predictions reveal SHD1-motif interactions across the Opisthokonta, demonstrating that the motif’s sequence, its interaction with SHD1, and its role in linking the polarisome to vesicle fusion are evolutionarily conserved.

## Results

### A conserved five-residue motif mediates binding of Msb3, Msb4, and Ste7 to Spa2

The Spa2 homology domain 1 of Spa2 (SHD1) is a conserved protein interaction module that is essential for the scaffold functions of Spa2 during bud growth and mating of yeast cells (Sheu et al., 1998; Lawson et al., 2022). A Split-Ubiquitin screen of SHD1CRU against an array of 530 N_ub_-fusion proteins, enriched for polar growth regulators, identified Msb3, Msb4 and Ste7 as interaction partners of SHD1 (Fig. 1A) (Hruby et al., 2011). Ste7 is the MAP kinase kinase in the kinase signaling pathway of the mating response of yeast cells (Courchesne et al., 1989; Teague et al., 1986). The GAP domains of Msb3, Msb4 and the kinase domain of Ste7 are preceded by intrinsically unstructured regions of 151, 59 and 187 residues in length respectively. All three regions were shown to be critical for the interaction with Spa2 (Sheu et al., 1998). Sequence analysis reveals a shared five-residue motif (V/I)-(I/L)-D-(L/A)-Y (from here on FRM) within the N-terminal regions of all three proteins (Fig. 1B). A similar motif is also found in the N-terminal regions of Mkk1 and Mkk2, MAP kinases of the cell wall integrity pathway, previously reported to interact with SHD1 (Fig. 1B) (Sheu et al., 1998).

**Figure 1:**
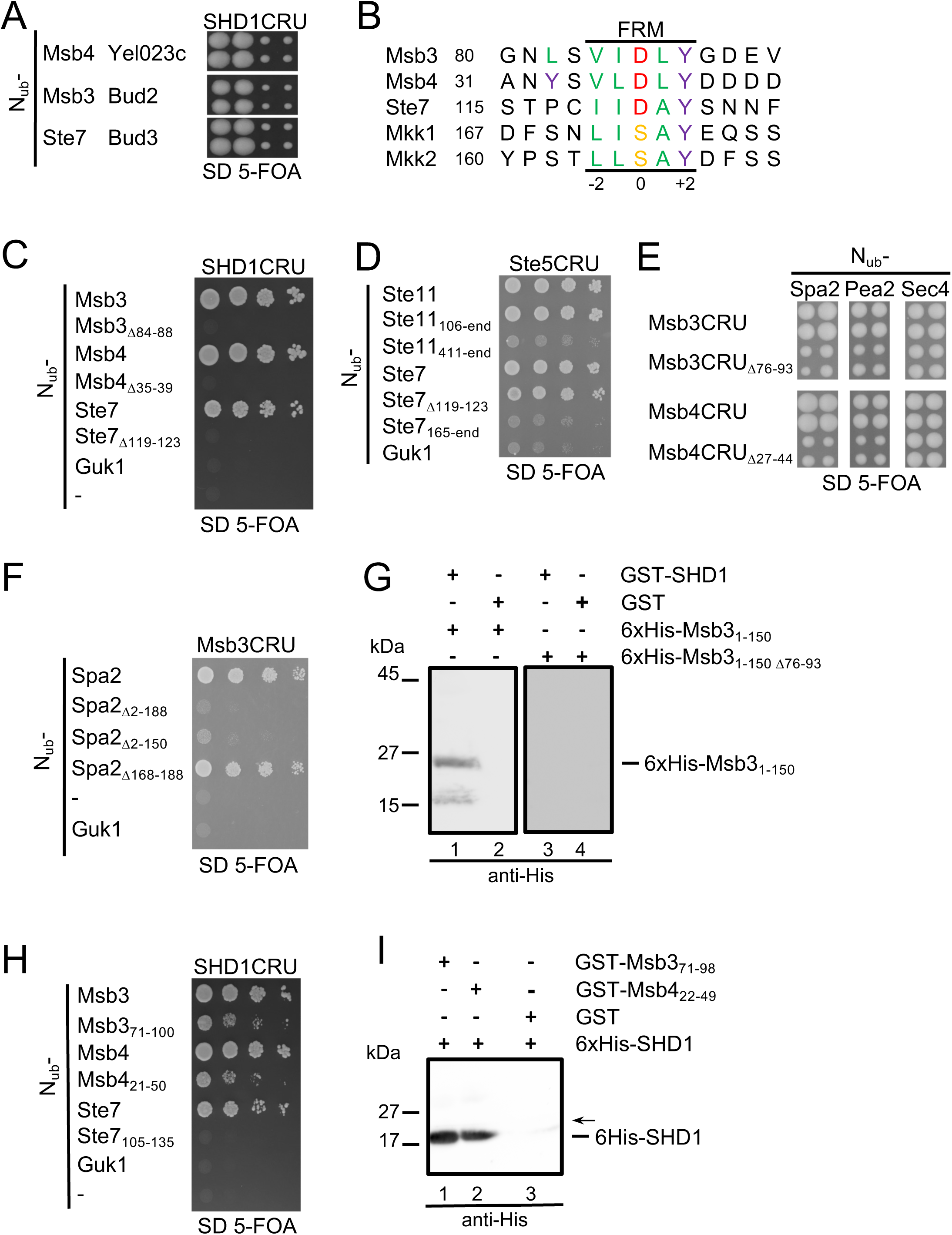
(A) Cut-outs of a Split-Ub array of diploid yeast cells each expressing SHD1CRU (CRU: C-terminal half of Ubiquitin (C_ub_)-R-Ura3) together with a different N_ub_ fusion (N_ub_: N-terminal half of Ubiquitin). Identities of the N_ub_ fusions are given next to the respective cut outs. Mated N_ub_- and C_ub_ expressing cells were arrayed as quadruplets on media containing 5-FOA. Colony growth indicates interaction between the fusion proteins. (B) Sequence alignment of putative binding sites of interaction partners of SHD1. FRM= Five residue motif. (C) Split-Ub assay of cells co-expressing SHD1CRU with N_ub_-fusions to SHD1-binding partners, their mutants lacking the respective FRM, or to Guk1 as negative control. Cells were grown to OD_600_=1 and 4 µl, or 4 µl of a 10-fold serial dilution were spotted on media containing 5-FOA and selecting for the presence of the N_ub_- and C_ub_ fusions. (D) As in (C) but with cells co-expressing Ste5CRU and N_ub_-fusions to Ste7, Ste11 and their mutants. (E) Split-Ub assay of cells co-expressing CRU-fusions to Msb3, Msb4 or their mutants lacking the FRM together with N_ub_-Spa2, N_ub_-Pea2, or N_ub_-Sec4. Shown are four independent transformants each, directly transferred from selective media onto 5-FOA media. (F) As in (C) but with cells co-expressing Msb3CRU together with N_ub_ fusions to Spa2 and mutant of Spa2 lacking the indicated regions of the protein. (G) Enriched 6xHis-Msb3_1-150_ (lanes 1-2) or 6xHis-Msb3_1-150Δ76-93_ (lanes 3, 4) were incubated with GST-SHD1 (lanes 1, 3) or GST-coupled beads (lanes 2, 4). Glutathione-eluates were separated by SDS-PAGE, transferred onto nitrocellulose, and stained with anti-His antibody (lanes 1-4) (Fig. S1A). (H) As in (C) but with cells co-expressing SHD1CRU together with N_ub_ fusions to Ms3, Msb4, Ste7 or to their peptides carrying the FRMs in central positions. (I) Enriched 6xHis-SHD1 (lanes 1-3) was incubated with bead-coupled GST-Msb3_71-89_, GST-Msb4_22-49_ or GST (lanes 1, 2, 3). Glutathione-eluates were separated by SDS-PAGE, transferred onto nitrocellulose, and stained with anti-His antibody (lanes 1-3) (Fig. S1B). The arrow indicates the position where the membrane was horizontally cut before antibody incubation to prevent rebinding of the detached 6xHis-SHD1to the immobilized GSTMsb3_71-89_ or GST-Msb4_22-49_. Split-Ub experiments were performed twice, pulldown analysis was performed four (1G) or three (1I) times.

Deletion of the FRM in N_ub_-Msb3_Δ84-88_, N_ub_-Msb4_Δ35-39_, and N_ub_-Ste7_Δ119-123_ abolished their binding to SHD1CRU (Fig. 1C). Ste7’s N-terminal region also binds the mating scaffold Ste5 (Fig. 1D). This interaction is unaffected by the FRM deletion, as N_ub_Ste7_Δ119-123_ retains similar affinity for Ste5CRU as wild-type N_ub_-Ste7 (Fig. 1D). The FRM does not influence Msb3 or Msb4 binding to their substrate, the RabGTPase Sec4, as CRU fusions of Msb3, Msb4, and their FRM-deficient mutants (Msb3_Δ76-93_CRU, Msb4_Δ27-44_CRU) bind N_ub_-Sec4 with equal affinity (Fig. 1E). Conversely, N_ub_ fusions to Spa2 and the polarisome subunit Pea2 preferentially interact with CRU fusions of the wild-type GAPs (Fig. 1E) (Valtz and Herskowitz, 1996).

N-terminal deletion analysis of N_ub_-fusions to Spa2 confirmed SHD1 as the sole FRM-binding site for Msb3CRU (Fig. 1F). In pull-down assays, GST-Msb3_1-150_ precipitated 6xHis-SHD1, whereas GST-Msb3_1-150Δ76-93_ lacking the FRM did not (Fig. 1G, S1A). Short FRM-containing regions of Msb3 (N_ub_-Msb3_71-100_) and Msb4 (N_ub_Msb4_21-50_), but not Ste7 (N_ub_-Ste7_100-135_), were sufficient for SHD1CRU interaction in the Split-Ubiquitin assay (Fig. 1H). Consistently, bacterially expressed GST-Msb3_71-98_ and GST-Msb4_22-49_ precipitated 6xHis-SHD1 *in vitro* (Fig. 1I, S1B).

### Alpha-Fold predicts the FRM-SHD1 complex with high confidence

Alpha-Fold predicts a compact globular fold for SHD1 where five helices create a groove lined by hydrophobic side chains (Fig. S1C)(Abramson et al., 2024). The rim of the groove exposes positively charged residues. The prediction is in good agreement with the solved structures of the SHD1s of the Spa2 homologues from *Neurospora crassa* and from mouse (GIT1) (Zheng et al., 2020; Zhu et al., 2020). The FRMs of Msb3, Msb4, Ste7 and Mkk1 reside in unstructured regions of the proteins. When confronted with the sequence of SHD1, AlphaFold converts these unstructured, FRM-containing regions into alpha helices and grafts them into the hydrophobic pocket of SHD1 (Fig. 2A, S2). The model of the complex predicts that Tyr at position +2 of the motif and the hydrophobic residues at position −2, −1 and to a lesser extent +1 contact the hydrophobic face of the SHD1 groove. Asp at position 0 of Msb3/4 and Ste7 is oriented toward K97 of SHD1 (Fig. 2A). The hydroxyl group of Y+2 is hydrogen-bonded to D112 of Spa2. The sequence alignment of the Msb3 homologues of different yeasts underscores the importance of the hydrophobic character of the residues at positions −2, −1, and the Tyrosine at +2 (Fig. 2B). To independently test the model of the complex, we exchanged each residue of the FRM of Msb3 individually against an Ala, and in addition the conserved Leu at position −4 against Tyr. Split-Ub analysis of interactions of the N_ub_-fusions showed that the exchanges D0A, and L-4Y outside of the motif do not measurably decrease the strength of the interaction between N_ub_-Msb3 and SHD1CRU whereas the Ala-exchanges at positions −2, −1 and +2 disrupt the complex, and the exchange at position +1A only slightly impairs the interaction (Fig. 2C). The replacement Y+2F strongly reduces complex stability (Fig. 2C). The low sensitivity of position +1 against the Ala-exchange is in line with the observation that the FRM of Ste7 displays an Ala and Msb3 of *Candida albicans* a Glu at this position (Fig. 1B, 2B). All Msb3 alleles were still predicted by AlphaFold to interact with SHD1. However, the scores of the interface predicted template modelling (ipTM) vary among the complexes and loosely correlate with the binding strength of the measured Split-Ub interactions. Mutations that impair the binding reduce the ipTM scores accordingly (Fig. 2C, lower panel). The Y+2A and the D0A exchanges had comparable effects on the complex stability of Msb4-SHD1 (Fig. 2D). This observation permitted us to generalize the findings on the FRM_Msb3_-SHD1 interaction to the motifs of the other Spa2 binding partners. The Leu at positions 48, 51, 59, 98, 101, and 109 of Spa2 are predicted to contribute to the hydrophobic patch of the binding pocket of SHD1 (Fig. 2E, lower panel). We exchanged positions 48, 51, 101 and 109 individually or in combination against Ala and measured the interactions of the mutants against the N_ub_-fusions of Msb4, Msb3, and Ste7. Each single residue exchange abolished the Split-Ub measured interaction between SHD1 and N_ub_-Ste7 or N_ub_-Msb3 (Fig. 2E). The dissolution of Split-Ubmeasured interaction between Msb4 and SDH1 required two simultaneous exchanges in the SHD1 domain (L48A and L51A; L101A and L109A) (Fig. 2E). Exchanging K97 against D abolished the interaction between SHD1_K97D_ and Msb3 but hardly affected the interaction between SHD1_K97D_ and Msb4 or Ste7 (Fig. 2E).

**Figure 2:**
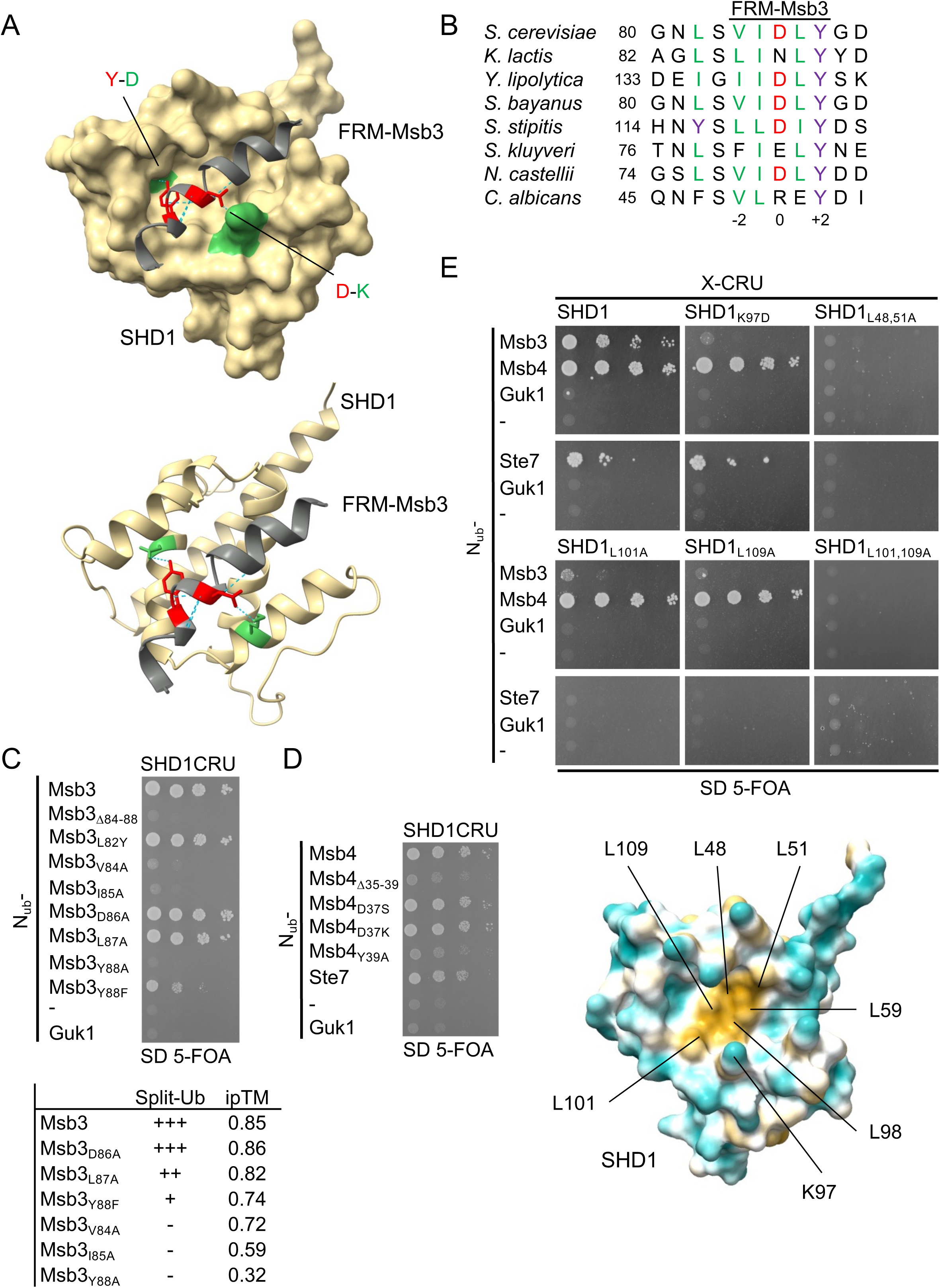
(A) Upper and lower panel: AlphaFold prediction of the Msb3-SHD1 complex. The FRM containing helix (grey) of Msb3 docks into the binding pocked of SHD1 (ochre). Y88 and D86 of Msb3 are indicated in red and D112 and K97 of Spa2 in green. The PAE scores of the predictions of the complexes between SHD1 and Msb3, Msb4, Ste7, and Mkk1 and the structures of the respective FRM-containing helices are shown in Fig. S2A-F. (B) Alignment of the sequences around the FRM of the Msb3 homologues from different yeast species. (C) Upper panel: Split-Ub analysis as in Fig.1C but with cells co-expressing SHD1CRU together with N_ub_ fusions of Msb3 or its mutants containing the indicated residue exchanges. Lower panel: Correlation between the ipTM scores of the predicted Msb3-SHD1 complexes carrying different residue exchanges in the FRM, and the estimated interaction strengths according to the Split-Ub assay. (D) Split-Ub analysis as in Fig.1C but with cells co-expressing Shd1CRU together with N_ub_ fusions of Msb4, mutants of Msb4 containing the indicated residue exchanges, and Ste7. (E) Upper panel: Split-Ub analysis as in Fig.1C but with cells co-expressing N_ub_ fusions to Msb3, Msb4 or Ste7 together with SHD1CRU or mutants of SHD1, carrying the indicated residue exchanges. Lower panel: Structural model of SHD1 by AlphaFold, highlighting the positions of the residues that form the hydrophobic patch of the binding groove. Split-Ub analysis was perfomed twice (2C, D), or four times (2E).

### Ste7, Msb3 and Msb4 compete for binding to Spa2

The similarity of the AlphaFold-structures and the comparable influences of the mutations within SHD1 on the interactions with its ligands strongly suggest but do not prove that Msb3, Msb4 and Ste7 compete for the same interface of Spa2. AlphaFold prediction shows that FRMs once fused to the N-terminus of Spa2 will fold back into the groove of the SHD1 and should consequently block any intermolecular binding of those proteins that use the same site on Spa2 (Fig. 3A). We devised a competition assay where we extended the N-terminus of the full length Spa2 with residues 22-81 of Msb4 (Msb4_22-81_-Spa2) (foldback competition assay, Fig. 3A). The N-terminal extension eliminated the binding of Msb4_22-81_-Spa2CRU with N_ub_-Msb3, and N_ub_Msb4 but did not affect the interactions with the N_ub_-fusions of the known Spa2binding partners Pea2, Epo1, and Aip5 (Fig. 3B) (Neller et al., 2015; Glomb et al., 2019). Deleting the FRM in the Msb4 extension (Msb4_22-81Δ35-39_-Spa2) recovered the interactions of Msb4_22-81Δ35-39_-Spa2CRU with N_ub_-Msb4 and N_ub_-Msb3 (Fig. 3B). The interaction between the full length Spa2CRU and N_ub_-Ste7 is not detected by the Split-Ub assay. To confirm that FRM_Ste7_ shares with FRM_Msb3/Msb4_ the same binding site on SHD1, we fused residues 105-165 of Ste7 in front of Spa2 (Ste7_105-162_Spa2CRU). As for Msb4_22-81_-Spa2, Ste7_105-162_ blocked the interaction between Ste7_105-162_-Spa2CRU and N_ub_-Msb3 and N_ub_-Msb4 (Fig. 3B). Alternatively, we extended the N-terminus of the isolated SHD1 by the unstructured FRM-containing regions of Msb3, Msb4, Ste7 and tested their influence on the interaction between the SHD1CRU and N_ub_-Msb3. The N-terminal extensions of Msb3/4 and Ste7 impaired the interaction between N_ub_-Msb3 and SHD1CRU whereas those extensions lacking a functional FRM did not (Fig. 3C).

**Figure 3.**
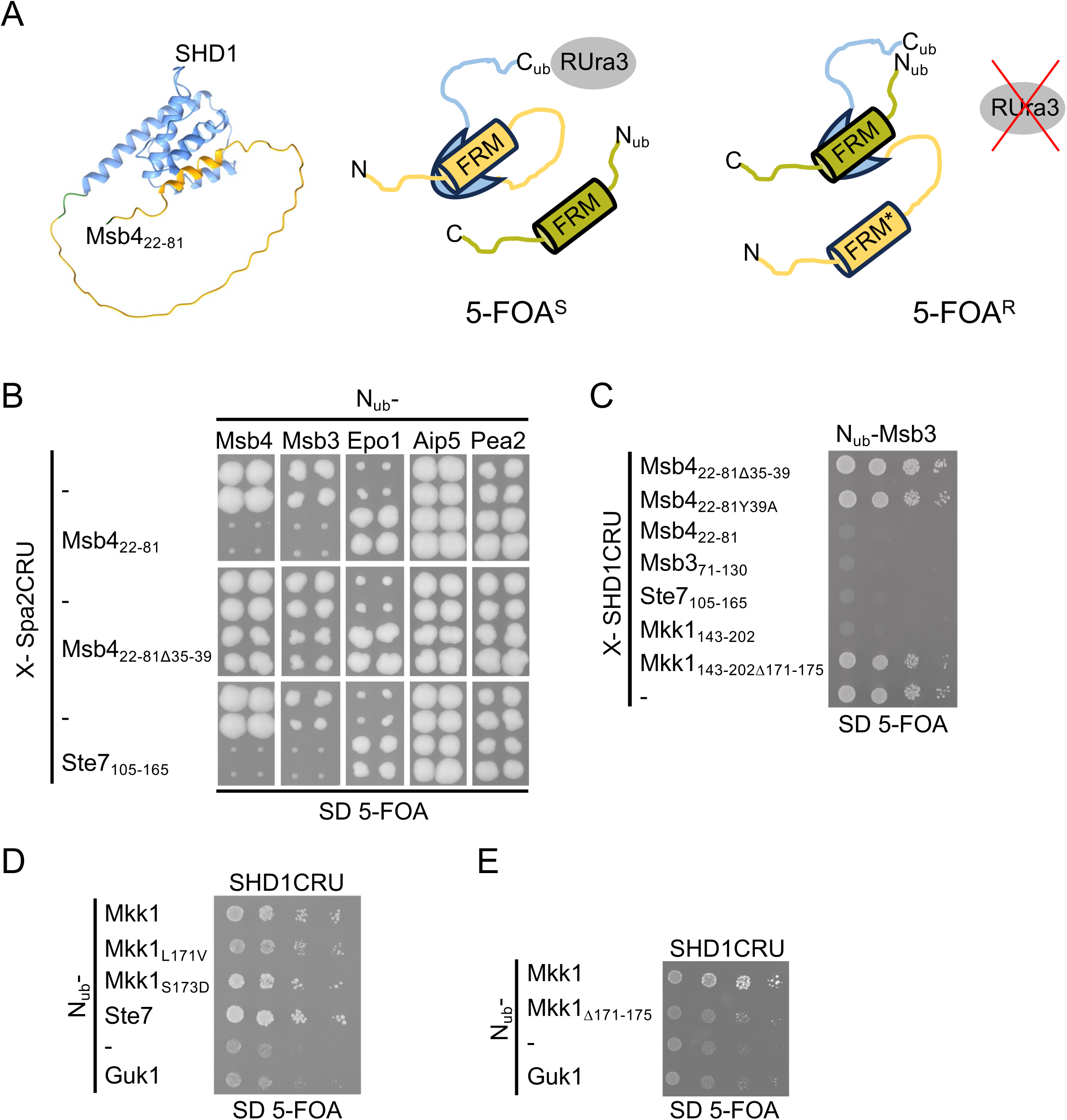
(A) A novel foldback competition assay shows that Mkk1 binds Spa2. Left panel: AlphaFold-predicted structure of SHD1(blue) extended at its N-terminus by residues 22-81 of Msb4 (ochre). Right panel: Foldback competition assay: Spa2 or its SHD1 domain (blue) are artificially extended at their N-termini by an FRM-containing amino acid stretch (ochre) and at their C-termini by the C_ub_RUra3 module (red). The FRM occupies intramolecularly the binding pocket of SHD1. As a consequence, an additionally expressed N_ub_-fusion to an FRM-containing protein (green) will not bind to SHD1, N_ub_ and C_ub_ will not reconstitute the native-like ubiquitin, and RUra3 will not be cleaved from C_ub_. The cells remain 5-FOA sensitive. Introducing an interactioninterfering mutations (FRM*) or deleting the FRM in the Spa2/SHD1 fusion protein will promote the intermolecular binding of the N_ub_-FRM fusion and the subsequent cleavage and degradation of the RUra3 module. The cell become 5-FOA resistant and grow in media containing 5-FOA. (B) Split-Ub assay as in Fig. 1E but with cells co-expressing Spa2CRU extended at its N-terminus by the N-terminal regions of Msb4, Msb4_Δ35-39_, or Ste7 together with N_ub_ fusions of the indicated proteins. (C) Split-Ub analysis as in Fig.1C but with cells co-expressing N_ub_-Msb3 together with SHD1CRU extended at its N-terminus by the FRM-containing regions of Msb4, Msb3, Ste7, or Mkk1. (D) Split-Ub analysis as in Fig.3D but with cells co-expressing SHD1CRU together with N_ub_ fusions of Mkk1, Mkk1_Δ171-175_, or its mutants containing the indicated residue exchanges on media containing 10mM methionine and 150 µm copper. (E) Split-Ub analysis as in (C) but with cells co-expressing SHD1CRU together with N_ub_ fusions of Mkk1, or Mkk1_Δ171-175_. Split-Ub analyses in (B)–(D) were performed twice.

### Mkk1 contains a FRM with a week affinity to SHD1

Sequence alignment reveals that Mkk1 possesses an FRM, yet N_ub_-Mkk1 was not identified as a binding partner of SHD1CRU in our initial screen (Mkk2 is absent from the N_ub_-fusion collection) (Fig. 1B). This could result from low affinity between SHD1 and Mkk1, low N_ub_-Mkk1 expression, or differing cellular localization of CRU- and N_ub_-fusion proteins. To address these factors, we employed the foldback competition assay, which is less sensitive to such variables (Fig. 3A, C). We fused residues 143– 202 of Mkk1 to SHD1CRU (Mkk1_143–202_SHD1CRU). Similar to the corresponding extensions of Msb3, Msb4 or Ste7, Mkk1_143–202_ blocked the interaction of Mkk1_143– 202_SHD1CRU with N_ub_-Msb3 (Fig. 3C). Deleting the FRM in Mkk1_143–202_SHD1CRU (Mkk1_160–204Δ171–175_SHD1CRU) restored this interaction (Fig. 3C). The foldback competition assay reliably detects FRM-SHD1 interactions even for Mkk1 that binds SHD1 with lower affinity than Msb3, Msb4, or Ste7 (see also Fig.6B). This finding aligns with AlphaFold predictions, which assign the Mkk1-SHD1 complex the lowest ipTM score among the four FRM-SHD1 complexes (Fig. S2; see also Fig. 7).

To confirm FRM_Mkk1_ binding to SHD1, we repeated the intermolecular Split-Ub assay with reduced SHD1CRU expression to enhance sensitivity. This revealed a weak but specific interaction between SHD1CRU and N_ub_-Mkk1 (Fig. 3D). Inspection of the FRM sequence in Mkk1 and Mkk2 identified a serine at position 0, unlike the more common aspartate in Msb3/4 and Ste7 (Fig. 1B). Substituting Ser0 with Asp slightly enhanced the interaction signal, whereas replacing Leu2 with the more prevalent Ile had no effect. Deleting the FRM in N_ub_-Mkk1 (N_ub_-Mkk1Δ171–175) reduced the interaction with SHD1CRU to background levels (Fig. 3E).

### The SHD1_Spa2_-FRM interaction targets the complex to sites of polar growth

Spa2 recruits Msb3 and Msb4 to the bud tip during bud growth, to the bud neck during mitosis and to the tip of mating projections (Tcheperegine et al., 2005; Lawson et al., 2022). Ste7-GFP localizes visibly only at the tip of mating projections (Maeder et al., 2007). To compare the influence of the FRM on the localization of each of the three proteins, we measured the cellular distributions of the respective GFP-fusions and their mutants upon exposure to mating hormone. Removing the FRMs from Msb3/4 and Ste7, or the SHD1 from Spa2 (Spa2_ΔSHD1_) abolishes the shmoo tip localization of Msb3-GFP, Msb4-GFP, and Ste7-GFP (Fig. 4A-D). In addition, tip and bud neck signals of the GFP-fusions to Msb3 and Msb4 in mitotic cells are lost or very much reduced when the FRMs of Msb3, Msb4 or the SHD1 of Spa2 are deleted (Fig. 4E).

**Figure 4.**
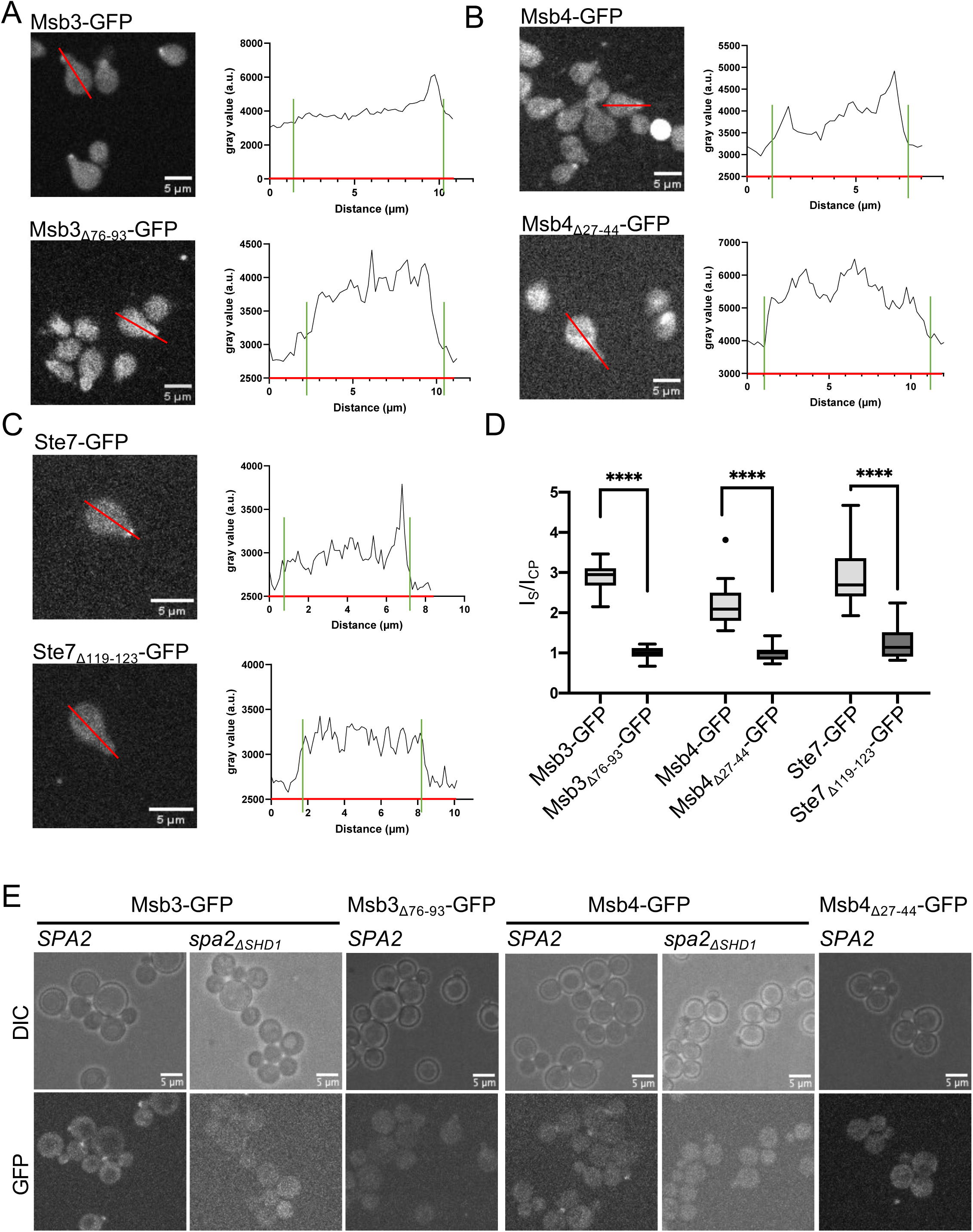
(A) Fluorescence microscopy of cells expressing Msb3-GFP (upper left panel) or Msb3_Δ76-93_-GFP (lower left panel) after treatment with mating factor. Red lines indicate sites of the fluorescence intensity measurements. Right panels: Fluorescence intensity profiles across the indicated cells. Green bars indicate the borders of the cell. Shmoo-tip ends at the right bar. (B) As in (A) but with cells expressing Msb4-GFP, or Msb4_Δ27-44_-GFP. (C) As in (A) but with cells expressing Ste7-GFP, or Ste7_Δ119-123_-GFP. (D) Shmoo-tip to cytosol fluorescence intensity ratios of cells expressing GFP fusions to Msb3 (n=10), Msb3_Δ76-93_ (n=10), Msb4 (n=10), Msb4_Δ27-44_ (n=10), Ste7 (n=10) or Ste7_Δ119-123_ (n=10) (Table S3). Whiskers of the box plots defined by the Tukey method. A-C, E representative images from at least two independent clones.

### The SHD1-FRM_Msb3/4_ complex directs vesicle fusion to the tip of the cell

The simultaneous loss of Msb3 and Msb4 induce the accumulation of post Golgi vesicles in the buds of *msb3Δmsb4Δ*-cells (Fig. 5A) (Gao et al., 2003; Tcheperegine et al., 2005). To investigate the influence of the Spa2-Msb3/4 interaction on the fusion of post-Golgi vesicles, we deleted the FRM of Msb3 in a *msb4Δ*-strain (*msb4Δ msb3_Δ76-93_*), the FRM of Msb4 in a *msb3Δ* strain (*msb3Δ msb4_Δ27-44_)*, or the SHD1 of Spa2 (*spa2_ΔSHD1_*). Electron microscopy of these cells did not reveal the characteristic accumulation of 50-70 nm vesicles that are prominently seen in *msb3Δmsb4Δ*-cells (Fig. 5A, B). Accordingly, no vesicle accumulation occurs in cells that lack the SHD1 of Spa2 (*spa2_Δ2-150_*), or lack Spa2 (*spa2Δ*) completely (Fig. 5A, B).

**Figure 5.**
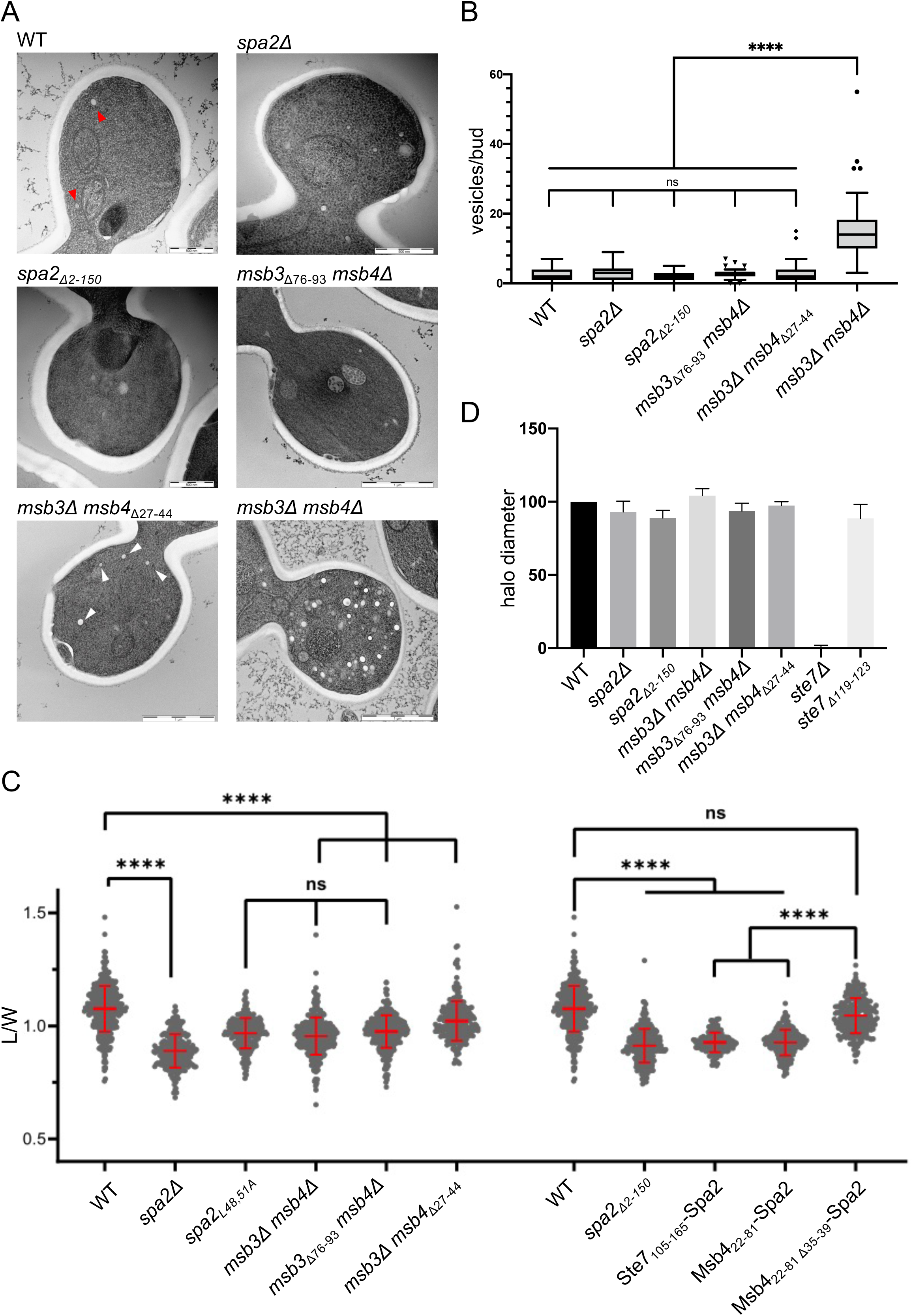
(A) Electron microscopical images of buds of cells of the indicated genotypes. Red arrowheads point at post-Golgi vesicles. (B) Quantification of 50-70 nm vesicles in 30 buds each of wt-, *spa2Δ, spa2_Δ2-150_*-*, msb3_Δ76-93_ msb4Δ*-*, msb3Δ msb4_Δ27-44_*-, and *msb3Δ msb4Δ-*cells. Whiskers of the box plots defined by the Tukey method. (C) Length/width ratios of buds of cells of wt (n=400), *spa2Δ* (n=240)*, spa2_L48,51_* (n=224)*, msb3Δ msb4Δ* (n=353)*, msb3_Δ76-93_ msb4Δ* (n=342)*, msb3Δ msb4_Δ27-44_* (n=291), *spa2_Δ2-150_* (n=264)*, ste7_105-165_spa2* (n=168)*, msb4_22-81_spa2* (n=149), *msb4_22-81Δ35-39_spa2* (n=233). The value for wt-cells is shown twice for clarity. (D) Diameters of zones of growth inhibition after central application of alpha-factor on a lawn of cells of the indicated genotypes (n=4; error bars=s.d.). EM images and quantifications (5A, B) from one clone each. Analysis in 5C from two independent clones each.

A tip-directed fusion of vesicles leads to ellipsoid bud shapes of the yeast, whereas random fusion leads to round bud shapes (Dünkler et al., 2021). The extent of tip-directed vesicle fusion can be approximated by the length to width ratio of the bud (L/W ratio). *spa2Δ-* and *msb3Δmsb4Δ-*cells have a very similar L/W ratio that is significantly smaller than those of wild type cells (Fig.5C). Cells expressing *msb3_Δ7693_* in *msb4Δ*-cells, or *msb4_Δ27-44_* in *msb3Δ*-cells display L/W ratios that are close to those of *spa2Δ-* or *msb3Δmsb4Δ*- cells. Deleting SHD1, or introducing the L48A L51A mutations in the hydrophobic pocket of SHD1 reduces the L/W ratio to a similar extent (Fig.5C). Accordingly, cells expressing Ste7_105-162_-Spa2, or Msb4_22-81_-Spa2, showed a significant reduced L/W ratio compared to cells expressing Msb4_22-81Δ35-39_Spa2 (Fig.5C). We conclude that the SHD1-Msb3/Msb4 interaction restricts vesicle fusion to the bud tip by recruiting the Sec4-GAP-activity to this site. Although the interaction between Ste7 and Spa2 is required to attach Ste7 to the shmoo tip (Fig. 4C, D), cells lacking this interaction arrest the cell cycle upon pheromone treatment like wildtype and not like *ste7Δ*-cells (Fig. 5D).

### The FRM of human β-PIX binds the SHD1 of Spa2

GIT1/2 are the mammalian homologues of Spa2 (Zhao et al., 2000). Their SHD1s are preceded by ankyrin repeats and an ARF GAP domain. The SHD1 domain of GIT1/2 binds to β-PIX and recruits the Cdc42-GEF to focal adhesions (Zhao et al., 2000). The solved crystal structure of the complex shows how the hydrophobic groove of SHD1_GIT1_ encloses a short helix of β-PIX (Zhu et al., 2020). Closer inspection of the β-PIX helix revealed the essential features of the yeast FRM (Fig. 6A). This observation encouraged us to test whether FRM_β-PIX_ binds to SHD1 of yeast. Probed on a blot by bacterially expressed 6xHis-SHD1_Spa2_, the overlay assay of the different GST-FRM fusion proteins demonstrates that FRM_β-PIX_ binds to SHD1 as strongly as FRM_Ste7_ but less strongly than FRM_Msb3/4_ (Fig. 6B). The N-terminal regions of Mkk1 or Mkk2 did not measurably interacted with 6xHis-SHD1 under these conditions (Fig. 6B).

**Figure 6.**
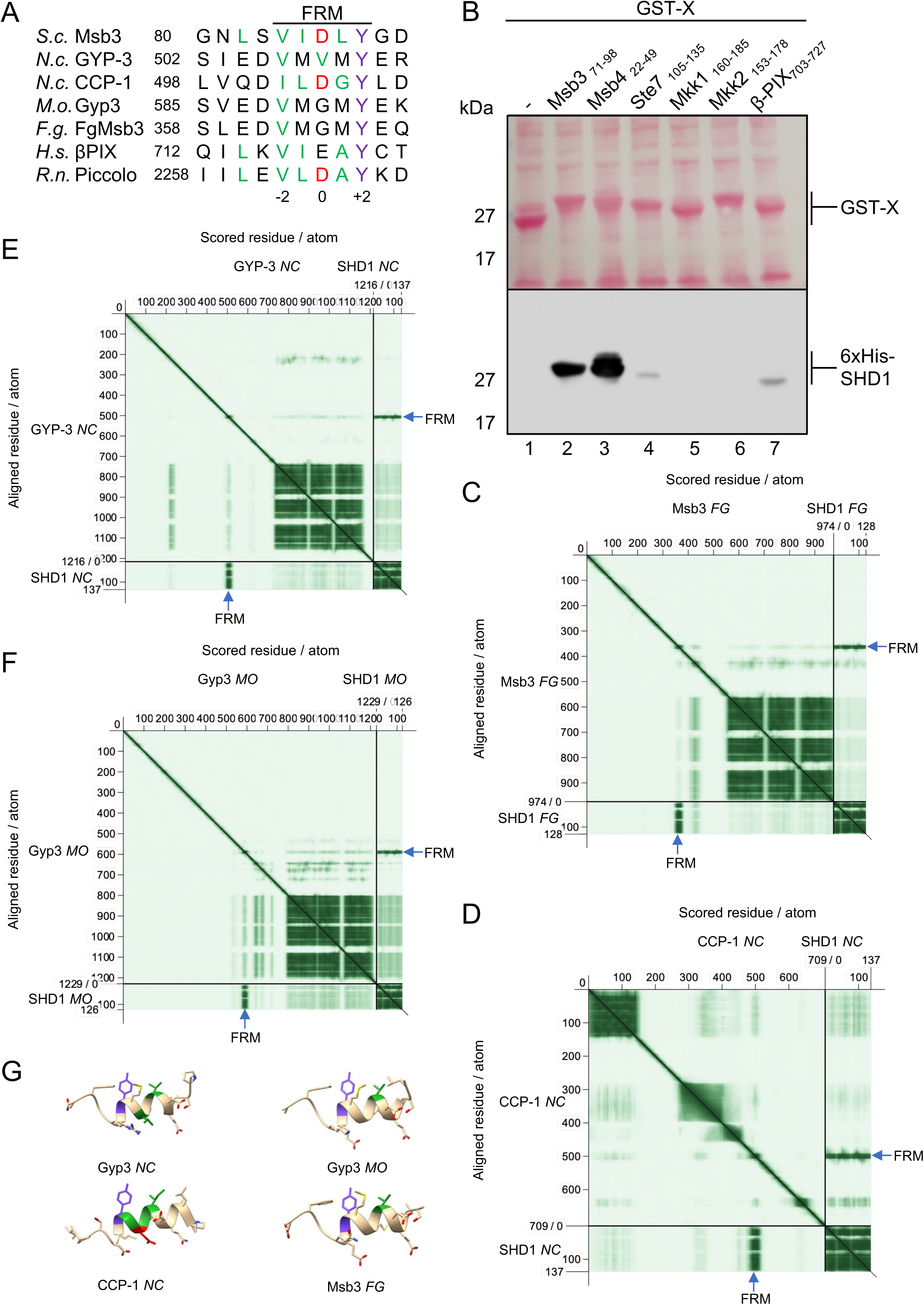
(A) Sequence alignment of the FRMs from different members of the *Ophistokonta*. (B) Extracts of *E.coli* cells expressing GST (lane 1), GST-Msb3_71-98_ (lane 2), GST-Msb4_22-49_ (lane 3), GST-Ste7_105-135_ (lane 4), GST-Mkk1_160-185_ (lane 5), GST-Mkk2_153178_ (lane 6), and GST-β-PIX _703-727_ (lane 7), were separated by SDS PAGE, transferred on nitrocellulose, stained with Ponceau S (upper panel) and then incubated with enriched 6xHis-SHD1 before being incubated with anti-His and peroxidase-coupled secondary antibody (lower panel). Experiment performed three times. (C-F) Predicted Aligned Error (PAE) matrices from the AlphaFold predictions of the complexes between the SHD1s of *Fusarium graminearum* (C), *Neurospora crassa* (D, E), and *Magnaporthe oryzae* (F) with the indicated binding partners. Blue arrows points to the sequences around the FRMs that display a high score of interaction with the SHD1. (G) Predicted structures of the FRM-containing sequences of the complexes of C-F.

### Predicting SHD1-FRM complexes in other members of the Opisthokonta

The similarity between the predicted SHD1-FRM complex of yeast and the GIT1-βPIX complex of mammals suggests, that SHD1s might generally bind sequence-conserved FRMs. Spa2 was discovered in polarisome-like structures of other yeasts and filamentous fungi (Knechtle et al., 2003; Jones and Sudbery, 2010). The architectures and compositions of these protein assemblies are not fully worked out and evidence for a physical interaction between Spa2 and Msb3/4 is only provided for the polarisome of *Fusarium graminearum* (FG), an ascomyceteous fungus that causes head blight in wheat (Zheng et al., 2021). The interaction between Spa2_FG_ and Msb3_FG_ were mapped to the SHD1 of Spa2_FG_ and to residues 168-399 in the supposedly unstructured region of Msb3_FG_ (Zheng et al., 2021). Supplied with the sequences of the SHD1 of Spa2_FG_ and the sequence of full-length Msb3_FG_, AlphaFold models with high confidence a short helix of Msb3_FG_ into the groove of Spa2_FG._ (Fig. 6C). The sequence of the helix matches the characteristic FRM of the budding yeast, and, by stretching from residues 362 to 365, falls exactly into the experimentally determined binding region of Msb3_FG_ (Fig. 6A, C, G).

By using AlphaFold as a tool to predict homologous SHD1-FRM-complexes, we explored whether RabGAPs are integral polarisome components in other species as well. The SHD1 of the Spa2 homologue from *Neurospora crassa* (NC) was shown to bind to the Calponin-like protein CCP-1 (Zheng et al. 2020). The binding region of Spa2_NC_ was assigned to the hydrophobic groove of SHD1, whereas the corresponding binding site on CCP-1 remained unspecified. AlphaFold predicts with high confidence a complex between both proteins. The interface of this complex is formed by the hydrophobic groove of SHD1_NC_ and a short helix from residue 497-507 of CCP-1 that matches the consensus of the FRM but carries a Gly instead of a hydrophobic residue in position +1 (Fig. 6A, D, G). Furthermore, indirect evidence suggest that Spa2_NC_ might also interact with Gyp3_NC_, the Msb3-homologue of *Neurospora crassa* (Callejas-Negrete and Castro-Longoria, 2019; Lichius et al., 2012). The predicted complex between SHD1_NC_ and full length Gyp3_NC_ confirms this suggestion. SHD1_NC_ captures a short helix with strong sequence similarity to the FRM. (Fig. 6A, E, G).

*Magnaporthe oryzae* (MO), is a pathogenic, filamentous fungi that causes rice blast. The polarsiome of *Magnaporthe oryzae* was shown to form at the tip of the hyphae (Li et al., 2014). A complex between Spa2_MO_ and Gyp3_MO_ was not described, but is predicted by AlphaFold with high confidence. Again, the helix of Gyp3_MO_ that fits into the groove of SHD1_MO_ is situated in the unstructured N-terminal region of Gyp3_MO_ and aligns well with the consensus sequence of the FRM (Fig. 6A, F, G). The mammalian GIT1 displays an internal Rab-GAP activity at its N-terminus thus freeing the SHD1 from its obligation to bind an external GAP. Instead, SHD1_GIT1_ binds besides β-PIX the scaffold protein Piccolo (Kim et al., 2003). The complex is part of the active zone below the presynaptic plasma membrane that might be considered to be functionally and structurally related to the cortical organization underlying the membrane of the yeast bud tip. The binding region of Piccolo from rat (RN) was experimentally localized within a fragment of 150 residues of the more than 5000 residue long protein (Kim et al., 2003). AlphaFold proposes a complex between SHD1_GIT1_ and a peptide within the binding fragment of Piccolo (Fig. S3A). The sequence of this peptide folds into a helix upon contact with the SHD1 binding cleft and matches the FRM consensus sequence (Fig. 6A, S3A-D).

### A proteome-wide search predicts Dse3 as novel interactor of Spa2

The relaxed stringency of the FRM’s consensus sequence increases the likelihood of its frequent, random occurrence within the yeast proteome. To distinguish functional motifs that directly interact with the SHD1 domain of Spa2 from these occurrences, we implemented the following strategy:

1. Identify all sequences conforming to the pattern (V/I)-(I/L)-(D/S)-(L/A)-Y.
2. Discard motifs not located in unfolded, or helical conformations.
3. Exclude non-cytosolic proteins.
4. Exclude proteins that, according to AlphaFold, do not bind the hydrophobic SHD1 groove through the motif’s helical conformation.
5. Exclude proteins as potential binding partners of SHD1 if the ipTM of the predicted complex does not scores equal to or higher than that of Mkk1.
6. Use the foldback competition assay to prove the motif-dependent binding of the candidate protein.

The strategy reduced the number of FRM containing proteins from 124 in Step1 to 95 in Step2, to nine in Step 4 (Fig. 7A). In Step 5, only the prediction of complexes between SHD1 and Msb3, Msb4, Ste7, or Dse3 sored equal or higher than the Mkk1SHD1 complex (Fig. 7A, Table S4). Similar to the motif-containing regions of Msb3, Msb4, Ste7 or Mkk1, Dse3_181-240_ inhibited binding of N_ub_-Msb3 to SHD1CRU in the Dse3_181-240_-SHD1CRU fusion protein (Step 6, Fig. 7B, C, Table S4). The inhibition was strictly dependent on the presence of the FRM as N_ub_-Msb3 interacted with Dse3_181-240Δ208-_212-SHD1CRU as strongly as with SHD1CRU alone (Fig 7C). The FRM-containing helix of Dse3, unlike the other yeast complexes, is predicted to adopt a flipped orientation within the binding groove. The Tyr residue at the +2 position of the motif forms a hydrogen bond with Asp62 of Spa2 (Fig. 7D, E).

**Figure 7.**
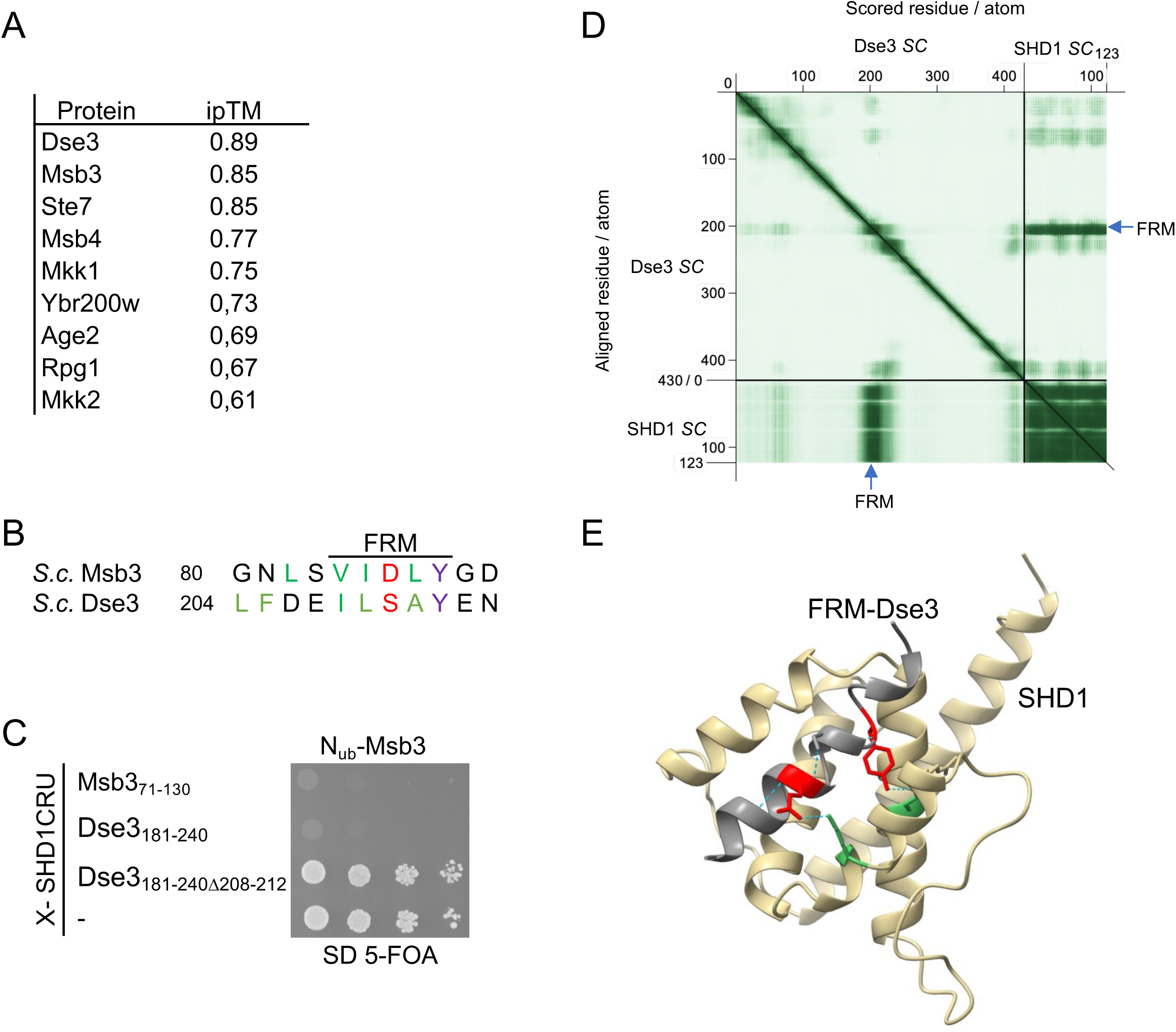
(A) List of the FRM-containing yeast proteins displaying the highest ipTM scores for the AlphaFold prediction of their complexes with the SHD1 of Spa2. (B) Sequence alignment of the FRMs of Msb3 and Dse3. (C) Foldback competition analysis as in Fig.1C but with cells co-expressing N_ub_-Msb3 together with SHD1CRU extended at its N-terminus by the FRM-containing regions of Msb3 (Msb3_71-130_-SHD1CRU), Dse3 (Dse3_181-240_-SHD1CRU), or its mutant lacking the FRM (Dse3_181-240Δ208-212_SHD1CRU) (performed twice, with two independent clones each). (D) Predicted Aligned Error (PAE) matrix from the AlphaFold prediction of the complex between the SHD1 of Spa2 and Dse3. (E) Structural model of the interaction between the FRMcontaining helix of Dse3 (grey) and SHD1 (ochre) derived from the AlphaFold prediction of the complex. E207 and Y212 of Dse3 highlighted in red; D62 and K97 of Spa2 highlighted in green. Split-Ub analysis (C) was performed twice.

## Discussion

The N-terminal domain of Spa2 binds to a conserved motif in Msb3, Msb4, Ste7, and Mkk1. Complex formation requires the initially unstructured motifs to adopt an alpha-helical conformation upon binding to the SHD1 domain. AlphaFold predictions, supported by mutational analysis of potential contact sites in Msb3-SHD1, Msb4SHD1, Ste7-SHD1 and Mkk1-SHD1 complexes, and *in vivo* competition assays among these FRMs, confirm this model. Independent validation comes from the solved structure of the mammalian GIT1 SHD1 domain bound to its partner, β-PIX (Zhu et al., 2020). The structures of the different complexes explain why the tyrosine at position +2 in the FRM is conserved across proteins and species. The tyrosine’s phenyl group forms hydrophobic contacts with the SHD1 groove, while its hydroxyl group forms a hydrogen bond with the carboxyl group of Asp112 in yeast Spa2 or Asp348 in mammalian GIT1, orienting the helix within the groove. Substituting tyrosine with phenylalanine, which lacks the hydroxyl group, significantly reduce the stabilities of both complexes (Fig. 2C and (Zhu et al., 2020)).

Surprisingly, the FRM orientation within the SHD1 hydrophobic groove is reversed in the predicted complexes from *Neurospora crassa*, *Magnaporthe oryzae*, *Fusarium graminearum*, and the complex between SHD1 and Dse3. Here, the tyrosine at position +2 forms a hydrogen bond with an aspartate corresponding to Asp62 in *Saccharomyces cerevisiae*. Both Asp112 and Asp62 are conserved in the SHD1 domains of yeast Spa2 and mammalian GIT1/2, occupying equivalent positions in SDR repeats 1 and 2, respectively, thus reflecting the internal symmetry of the SHD1 fold. Using Asp112 or Asp62 as the hydrogen bond acceptor requires reshuffling hydrogen bonds within SHD1 and reversing the helix orientation in the groove. Further experimental validation is needed to distinguish between these alternative arrangements of the helices in the binding groove of SHD1.

Our findings indicate that an FRM in an unstructured protein region strongly suggests binding to a member of the Spa2/GIT1 family of cortical scaffold proteins. Using AlphaFold to identify potential binding sites, rather than relying solely on sequence alignments, enables detection of sites with partial matches to a not fully defined consensus sequence, and checks in addition the spatial accessibility of the sequence at the same time. However, the weak correlation between interaction strength (measured by the Split-Ub assay) and ipTM scores in complexes with single-residue substitutions underscores the limitations of AlphaFold for these applications. A low ipTM score may still reflect an FRM with functionally relevant, though weaker affinity. If the strength of interactions affects the accuracy of complex predictions, establishing computational criteria to differentiate non-functional FRMs from those engaged in weak but functional interactions would be highly beneficial. This is especially crucial when translating insights from protein interaction networks in model organisms to species less amenable to experimental study.

Our assays demonstrate that Msb4, Msb3, Ste7, and Mkk1 bind Spa2 with decreasing affinity. These differences are physiologically relevant for interactions involving Spa2 with Msb3, Msb4, and Ste7. The stronger binding of Msb3 and Msb4 ensures their localization to the cell tip and bud neck during mitotic growth and mating, whereas Ste7’s weaker affinity for SHD1 likely requires additional interactions with proteins like Ste5, which co-assemble at the cell tip exclusively during mating to stabilize Ste7 alongside Spa2 (Choi et al., 1994;,Maeder et al., 2007). Similarly, Mkk1’s polar distribution depends on Spa2, despite its interaction with SHD1 being near the detection limit of our assays (Van Drogen and Peter, 2002). Mkk1 stabilization at these sites may require Slt2, the MAP kinase in this pathway, which binds to both Mkk1 and Spa2 (Hruby et al., 2011; Breitkreutz et al., 2010).

Our homology search for functional FRMs in the yeast proteome identified Dse3 as a novel FRM-containing protein interacting with Spa2. *DSE3* is specifically transcribed in daughter cells (Colman-Lerner et al., 2001). Nothing is known about the function of the expressed protein. The functional relevance of its interaction with Spa2 thus remains unclear and requires further exploration. The fold-back competition assay’s simplicity, sensitivity, and reliability make it an effective new tool for analyzing SLiMs, complementing existing methods (Subbanna et al., 2025). It enables efficient screening of previously overlooked candidates, such as Age2, Rpg1, Ybr200w-A, and Mkk2, for direct FRM-dependent interactions with Spa2. Additionally, it supports revisiting homology searches with less stringent FRM consensus sequences.

## Material and Methods

### Growth conditions, cultivation of yeast strains, and genetic methods

All yeast strains were derivatives of JD47, a segregant from a cross of the strains YPH500 and BBY45. Yeast strains were cultivated in synthetic defined (SD) or yeast extract peptone dextrose (YPD) media at the indicated temperatures. Media preparation followed standard protocols (Glomb et al., 2020). SD medium for Split-Ub assays contained in addition 1 mg/ml 5-fluoro-orotic acid (5-FOA; Formedium). Gene deletions and promoter replacements by *PMET17* were performed by homologous integration of the cassettes derived by PCR from the plasmids pFA6a-hphNT1, pFA6a- natNT2, pFA6a-kanMX6, pFA6a-CmLEU2, or pYM-N35 (Bähler et al., 1998; Janke et al., 2004). *E. coli* XL1 blue cells were used for plasmid amplification and grown at 37°C in lysogeny broth (LB) medium containing antibiotics. Proteins were expressed in the *E. coli* strain BL21 DE3, grown in LB or super broth (SB) medium at 37°C.

### Generation of plasmids and yeast strains

Detailed lists of all yeast strains and plasmids used in this study are provided in Table S1 and S2, sequence information on oligos is available upon request. Genomic deletions and modifications of the SHD1 or FRM motifs were generated by CRISPR/Cas9 manipulation using plasmid pML104 or pML107 containing specific 20mer guide RNA sequences, and template oligonucleotides for exchanging the information on the genomic DNA (Laughery et al., 2015). The correct insertions of PCR fragments and mutations were verified by PCR amplification and sequencing.

Gene deletions were obtained by replacing the ORF through single-step homologous recombination with an antibiotic resistance cassette derived by PCR from the plasmids pFA6a-hphNT1, pFA6a- natNT2, pFA6a-kanMX6, pFA6a-CmLEU2, or pYM-N3(Janke et al., 2004) (Bähler et al., 1998). Genomic gene fusions to the GFP- or CRU-modules were obtained as described (Müller et al., 2024). The fusion of *GFP* or *CRU* to *SPA2*, *MSB3*, MSB4, Ste7, Mkk1, and derivates of those genes with deleted or modified FRM or SHD1 regions were constructed by PCR amplification of the respective C-terminal ORFs without stop codon from genomic DNA. The obtained DNA fragments were cloned via *Eag*I and *Sal*I restriction sites in front of the *CRU* or *GFP* module on pRS303, pRS304, or pRS306 vectors (Wittke et al., 1999). The plasmids were linearized using a single restriction site within the C-terminal genomic DNA sequence and transformed into yeast. Successful integration was verified by PCR of single yeast colonies with diagnostic primer combinations using a forward primer annealing in the target ORF but upstream of the linearization site, and a reverse primer annealing in the C-terminal module.

Fusions between the N-terminal regions of Msb4, or Ste7, and full length Spa2 were generated *via* CRISPR/Cas9-mediated insertion of a PCR-fragment encoding the respective regions of *MSB4* or *STE7* flanked by 45bp identical to the region upstream of the start codon of *SPA2* and the 5’-end of its open reading frame. The plasmid-borne fusions between SHD1 and the N-terminal regions of Msb4, Ste7, Mkk1 or Msb3 were obtained by integrating the PCR products of the respective regions in front of SHD1 in the pMET17-Spa2(1-124)CRU313 vector.

Fragments of *SPA2, Msb3 or Msb4* were expressed as GST- or 6xHis- fusions in *E.coli* strains BL21 or BL21 Gold. Fragments for the GST-fusions were amplified from yeast genomic DNA using primers containing *Bam*HI/*Eco*RI restriction sites. The PCR products were fused in-frame behind GST on a pGex2T vector (GE Healthcare, Buckinghamshire, UK). 6xHis-tagged fragments were amplified from genomic DNA of wild type yeast or the strain lacking residues 76-93 of Msb3 using primers containing *Sfi*I restriction sites. The products were inserted into the pES plasmid, downstream and in frame of a 6xHis-tag. GST-FRM fusions of MSB3, MSB4, STE7, MKK1, MKK2 and β-PIX were obtained by inserting double stranded oligonucleotides harboring the sequences of the FRMs and containing matching *Sal*I/EcoRI restriction sites at their ends in-frame behind the ORF of GST on a pGEX2T vector.

### *In vivo* Split-Ub interaction analysis

Large-scale Split-Ub assays were performed as described (Hruby et al., 2011). A library of 540 different α-strains, each expressing a different N_ub_ fusion, was mated with a *PMET17SHD1*-CRU–expressing a-strain. Diploids were transferred as independent quadruplets on SD media lacking methionine and containing 1 mg/ml 5FOA, and different concentrations of copper sulfate to adjust the expression of the N_ub_ fusions. For small-scale interaction analysis, a- and *α-*strains expressing N_ub_ or C_ub_ fusion constructs were mated. The diploid cells were spotted onto SD-FOA medium in four 10-fold serial dilutions starting from OD_600_ = 1. Growth was recorded at 30°C every day for 2–5 d.

### Binding assay

All incubation steps were carried out under rotation at 4°C. GST or GST-tagged proteins were immobilized from *E. coli* extracts on 50 μl Glutathione–Sepharose beads in PBS (GE Healthcare). After incubation for 1 hour at 4°C, with either *E. coli* extracts or purified proteins, the beads were washed three times, the bound material was eluted with GST elution buffer (50 mM Tris, 20 mM reduced glutathione), and subjected to SDS-PAGE followed by Ponceau S staining and western blot analysis using anti-His (H1029), or anti-GST (G1160) antibodies (Sigma-Aldrich). To prevent re-binding of detached 6×His-SDH1 to immobilized GST fusions during antibody incubation (Fig.1I), Ponceau-stained membranes were cut horizontally between the ∼16 kDa and ∼30 kDa regions immediately after transfer. The resulting strips were processed in parallel in separate trays with identical solutions and reassembled for chemiluminescent detection.

### Far-western blot

Enriched GST and GST-FRM-fusions were transferred onto nitrocellulose after SDSPAGE. The membrane was blocked for 1 h in 20mM Tris, 150 NaCl, 0.1% Tween 20 (TBST) 2% milk powder (w/v), washed for 5 min in 10 ml TBST, and incubated for 1 h at 4°C with enriched 6xHis-SHD1. After washing three times for 5 min with 10 ml TBST, the membrane was sequentially incubated at room temperature for 1 h with anti-His (H1029) and conjugated anti-mouse antibodies (A4416; Sigma-Aldrich) (Wu et al., 2007). The source data of the western and far-western blots are shown in Fig. S4.

### Halo Assay

Exponentially growing yeast strains were diluted to OD_600_ = 0.5 and 300 μL of the suspension were plated evenly on SD-full agar plates. Round filter papers with a diameter of 5 mm were placed on the cell lawn and 10 μL of alpha-factor was applied to the filter papers in serial dilutions of 10 μg, 5 μg, 2.5 μg, and 1.25 μg. Plates were incubated overnight (o/n) at 30°C. Images were taken on day one and two. Sensitivity to mating factor was determined by measuring the halo of growth inhibition. All strains were measured four times, and the values were normalised to the wild type.

### Fluorescence microscopy

Yeast cultures were grown overnight in SD medium, diluted in 3–4 ml fresh SD medium, and grown for 2–3 h at 30°C to mid-log phase. Microscopic observations were performed with the Axio Observer spinning disc confocal microscope (Zeiss) equipped with an Evolve512 electron-multiplying charge-coupled device camera (Photometrics), a Plan-Apochromat 63×/1.4 oil differential interference contrast (DIC) objective, and a 100×/1.4 oil DIC objective. Fluorescence was excited with 488- diode lasers (Zeiss) and detected with a high-efficiency filter set 38 (GFP). Time-lapse experiments were carried out at RT over the course of 4h. Operations were performed with the ZEN2.6 (2012) software package (Zeiss). Exposure times were adjusted to the respective GFP -labeled proteins to reduce bleaching and phototoxicity (Müller et al., 2024).

Standard time-lapse experiments were carried out with the 63× objective in a Sarstedt 1-well on cover glass II incubation chamber. 100 μl culture was pipetted onto the glass bottom of the chamber and covered with a slice of standard solid SD medium. Images from a series of 14 z-slices over a total range of 2.43 µm were taken in intervals of 3 min. Special glass slides with a ground indentation in the centre were used for monitoring shmoo formation of alpha-factor-treated yeast cells (custom-made by the University’s glassblowing workshop). α-mating factor was added to a concentration of 10 μg/mL to 500 μL in 2x SD full media, mixed 1:1 with 3.4% agarose (heated until liquid), and immediately poured into the indentation. A coverslip was placed on the top to remove excess agarose and to match the surface of the glass. After cooling, the coverslip was removed, 3.6 μL of cell suspension were applied, covered with a new coverslip and sealed with parafilm to prevent evaporation and drying of the agarose patch. The slide was then incubated at 30°C for 15 minutes to allow the cells to settle.

### Analysis of microscopic data

Image analysis and signal quantifications were carried out with the open source software platform Fiji (ImageJ) (Schneider et al., 2012; Schindelin et al., 2012). Plot profiles of fluorescence intensities were performed in FIJI to detect differences in GFP distributions across the cell. Circular regions of interest (ROI) of the same size were placed at the shmoo tips of cells to quantify the GFP intensity relative to signal of the cell body. The background signal (I[background]) was measured with ROIs placed outside but in proximity of the cell and subtracted from the intracellular signals. The relative GFP intensities at the shmoo tip were then calculated: 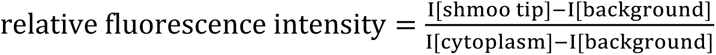

### L/W ratio determination

Yeast strains were grown overnight in SD medium and diluted in YPD to measure cell- and bud sizes. Images were taken on the Zeiss Observer Z1 using the Axio camera at 100x magnification with 14 z-layers over 3 μm in the DIC channel. Cells were randomly selected, and strain identities were blinded to avoid bias. The images were analysed in ImageJ (FIJI) after z-projection. To determine the bud morphology, the length (L) and width (W) of the buds were measured. The region of interest (ROI) for length was chosen along the axis of growth, and the ROI for width was set perpendicular to the length at the widest point of the bud. Only non-blurred, living cells with visibly attached buds were measured and strain names were kept blinded for evaluation. All values were transferred to Excel, sorted by length and the L/W ratio was calculated. Each strain was measured in triplicate, with a minimum of 150 cells measured. Buds smaller than 1.5 μm were excluded from the analysis.

### Transmission electron microscopy

Exponentially grown yeast cells were harvested and vitrified through high-pressure freezing on carbon coated sapphire discs in a Wohlwend HPF compact 01 highpressure freezer. Freeze-substitution was performed with 0.2% (v/v) osmium tetroxide, 0.1% (w/v) uranyl acetate and 5% (v/v) H_2_O in acetone by gradually increasing the temperature over 17 hours from −90 °C to 0 °C. Temperature was kept at 0 °C for one hour and then increased within one hour to room temperature. Samples were washed with acetone, followed by embedding in Epon 812 in four consecutive steps at room temperature: 33% Epon 812/67% acetone for 1 hour, followed by 67% Epon 812/33% acetone for 3 hours, 90% Epon 812/10% acetone overnight, and pure Epon 812 overnight (v/v). Samples were transferred into fresh Epon 812 and polymerized at 60 °C for 72 hours. Embedded cells were sectioned in ∼70 nm steps with a Leica EM UC7 ultramicrotome. Sections were captured on a formvar film on glow-discharged copper grids. Images were acquired with a 120 kV Jeol JEM-1400 Transmission electron microscope and a Veleta CCD camera (Olympus).

### Statistical evaluation

All statistical evaluations were based on data sets of at least three independent experiments. Depending on the result of the normality test, parametric (ANOVA) or non-parametric (Kruskal-Wallis) tests were chosen. The Kruskal-Wallis test was followed by a Dunn’s Test for multiple comparisons. The p values and the sample size (n) are given in the caption or in the supplementary data. Unless otherwise stated, the red bars represent the mean and standard deviation (s.d.) of each data set, the p values are given in GP standard values.

### Structure prediction

Structure prediction was performed with the AlphaFold 3 (Abramson et al., 2024). The models were ranked by their pTM/ipTM scores, and the model with the highest score is shown. The PAE scores were obtained from the AlphaFold3 data by the PAE viewer tool (Elfmann and Stülke, 2023). Processing and visualization of the predicted models were performed with ChimeraX (Meng et al., 2023).

### Proteome-Wide Motif Search

To identify amino acid sequences in Saccharomyces cerevisiae matching the motif (V/I)-(I/L)-(D/S)-(L/A)-Y, we analyzed the proteome-wide structure prediction dataset (n = 6,039) from the AlphaFold Protein Structure Database (version 4; available at: https://ftp.ebi.ac.uk/pub/databases/alphafold/v4/UP000002311_559292_YEAST_v4.t ar). A custom Bash script iteratively executed PyMOL (http://www.pymol.org/pymol.) (Schrödinger and DeLano, 2020) to scan each protein structure for the motif and document the secondary structure assignments of the corresponding five residues. Protein identifiers containing the motif were compiled into a comprehensive list of all matches. A separate list was generated for matches where the motif was exclusively located in residues annotated by PyMOL as helical or loop regions (including cases with both annotations).

### Use of artificial intelligence tools

Grok3 was used during manuscript preparation to provide suggestions for translation, grammar, and stylistic improvements.

## Data and Resource Availability

All strains and plasmids are available upon request.

## Acknowledgement

We thank C.M. Pfeiffer for helpful advice.

## Author contributions

Conceptualization: N. Johnsson. Investigation: L. Bareis, A. Siewert, S. Timmermann, N. Schmid, T. Bergner, Benjamin Grupp. Writing - original draft: N.

Johnsson; Writing - review & editing: N. Johnsson, L. Bareis. Supervision: N. Johnsson, C Read. Funding acquisition: N. Johnsson.

---

**Figure S1.**
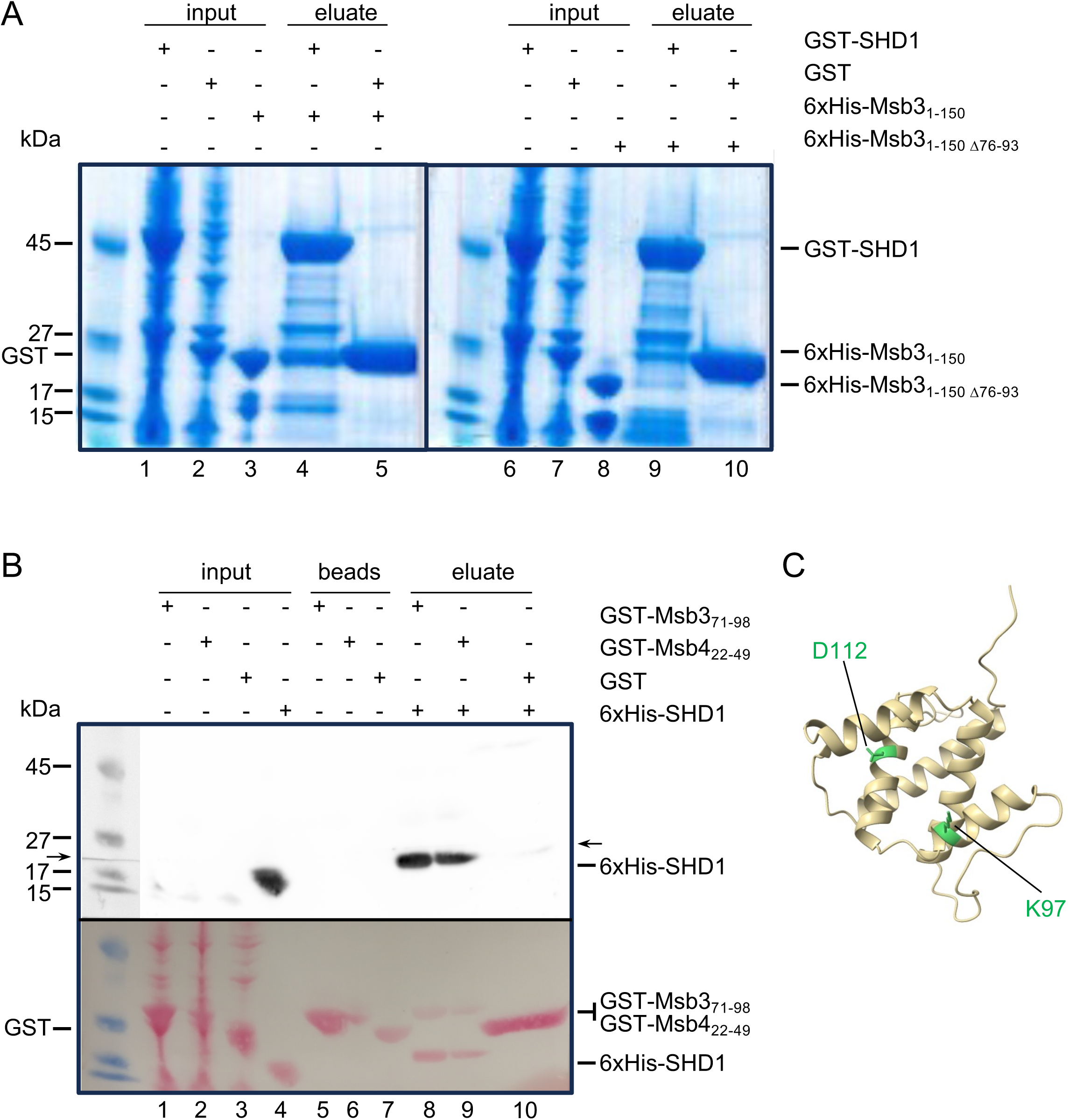
(A) Inputs for Fig.1 G. Coomassie staining of SDS-PAGE of extracts of *E.coli* cells expressing GST-SHD1 (lanes 1, 6) or GST (lanes 5, 10), the enriched 6xHis-Msb3_1-150_ (lane 3), or 6xHis-Msb3_1-150Δ76-93_ (lanes 8) and the eluates of GST-SHD1- (lane 4, 9) or GST- (lanes 5, 10) coupled beads after being incubated with the enriched 6xHis-Msb3_1-150_ (lanes 4, 5), or 6xHis-Msb3_1-150Δ76-93_ (lanes 9, 10). (B) Full analysis of the GST pull-down experiment shown in Figure 1I. Lanes 1–4: input; lanes 5–7: SDS eluate after glutathione eluate; lanes 8–10: glutathione eluate. Lanes 8–10 are identical to lanes 1–3 in Figure 1I. Upper panel: anti-His immunoblot. Arrow indicates position of the cut. Lower panel: Ponceau S stain of the same membrane before antibody incubation. (C) AlphaFold structure prediction of SHD1 from yeast.

**Figure S2.**
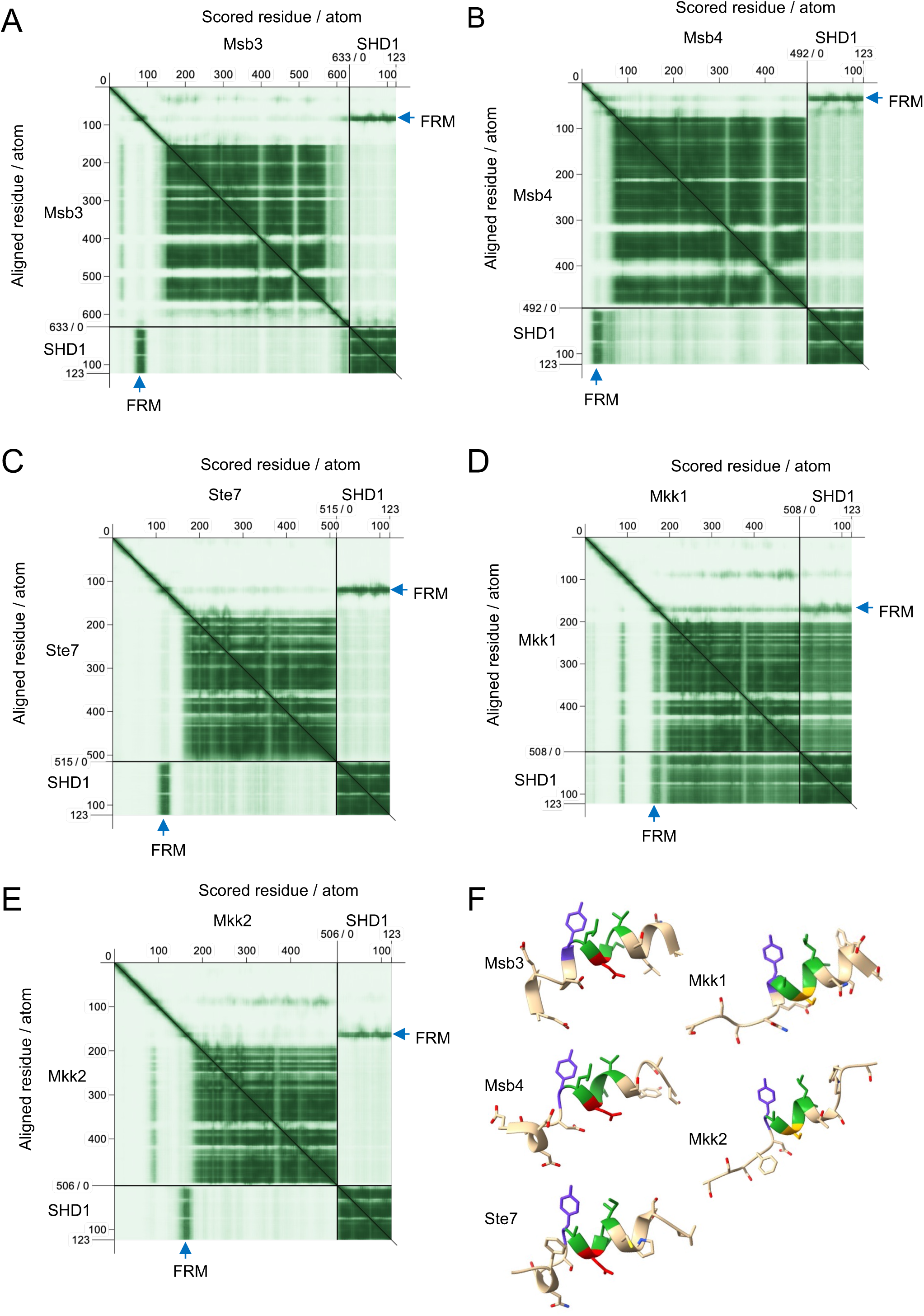
(A-E) Predicted Aligned Error (PAE) matrices from the AlphaFold predictions of the complexes between the SHD1 of budding yeast and the full length Msb3 (A), Msb4 (B), Ste7 (C), Mkk1 (D) and Mkk2 (E). Blue arrows point to the sequences around the FRMs. Dark green indicates a high probability score of interaction with the SHD1. (G) Predicted structures of the FRM-containing sequences of the complexes of A-E.

**Figure S3.**
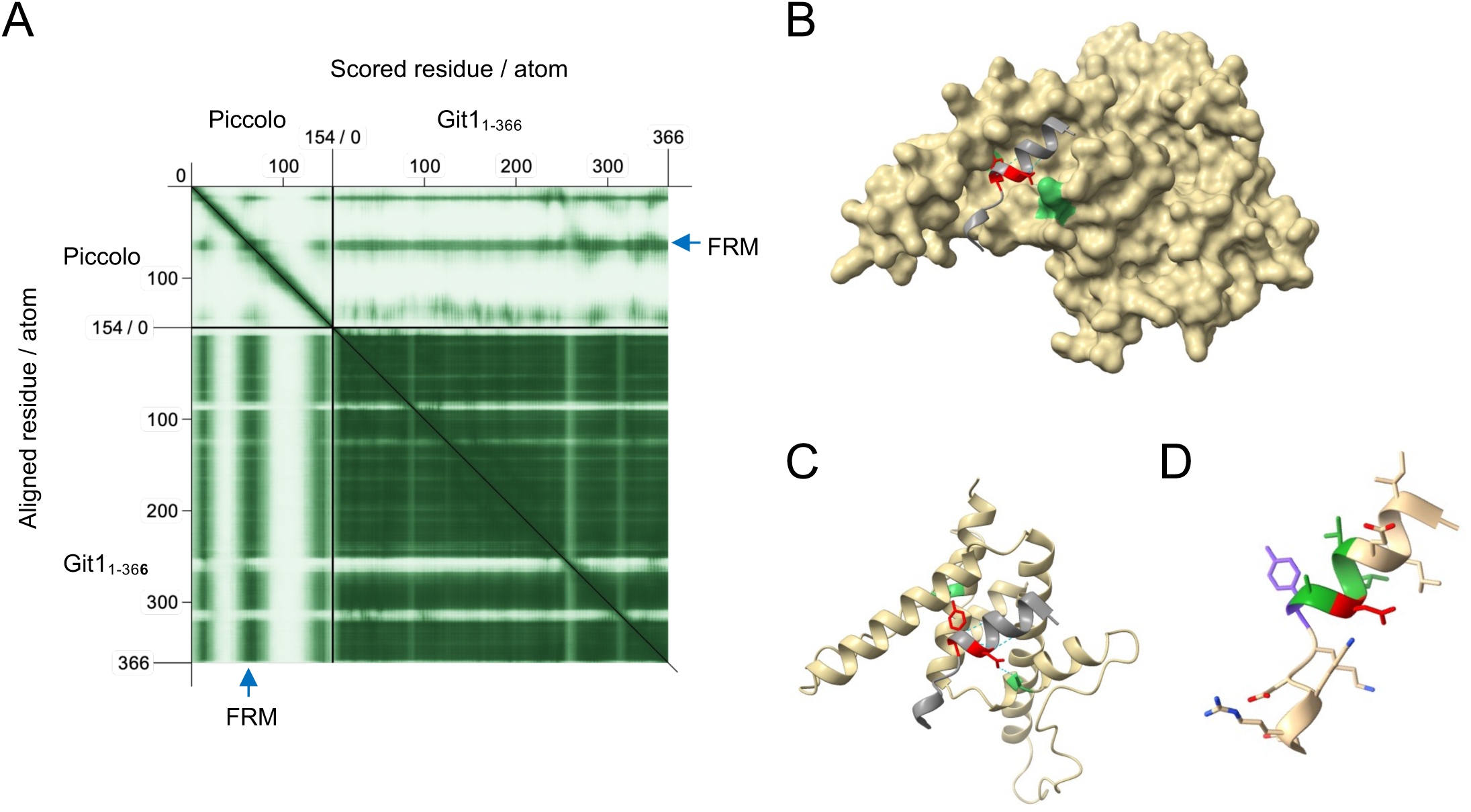
(A) Predicted Aligned Error (PAE) matrices from the AlphaFold predictions of the complexes between the GIT1_1-366_ of rat and a fragment of Piccolo. Blue arrows points to the sequences around the FRMs that display a high score of interaction with the SHD1 of GIT1. (B) AlphaFold structure of the complex between GIT_1-366_ and the FRM of Piccolo indicated in grey and highlighting the Y and D at the central positions. (C) Structure of the complex between the SHD1 of GIT1 and the FRM of Piccolo. (D) Structure of the helix containing the FRM of Piccolo.

**Figure S4.**
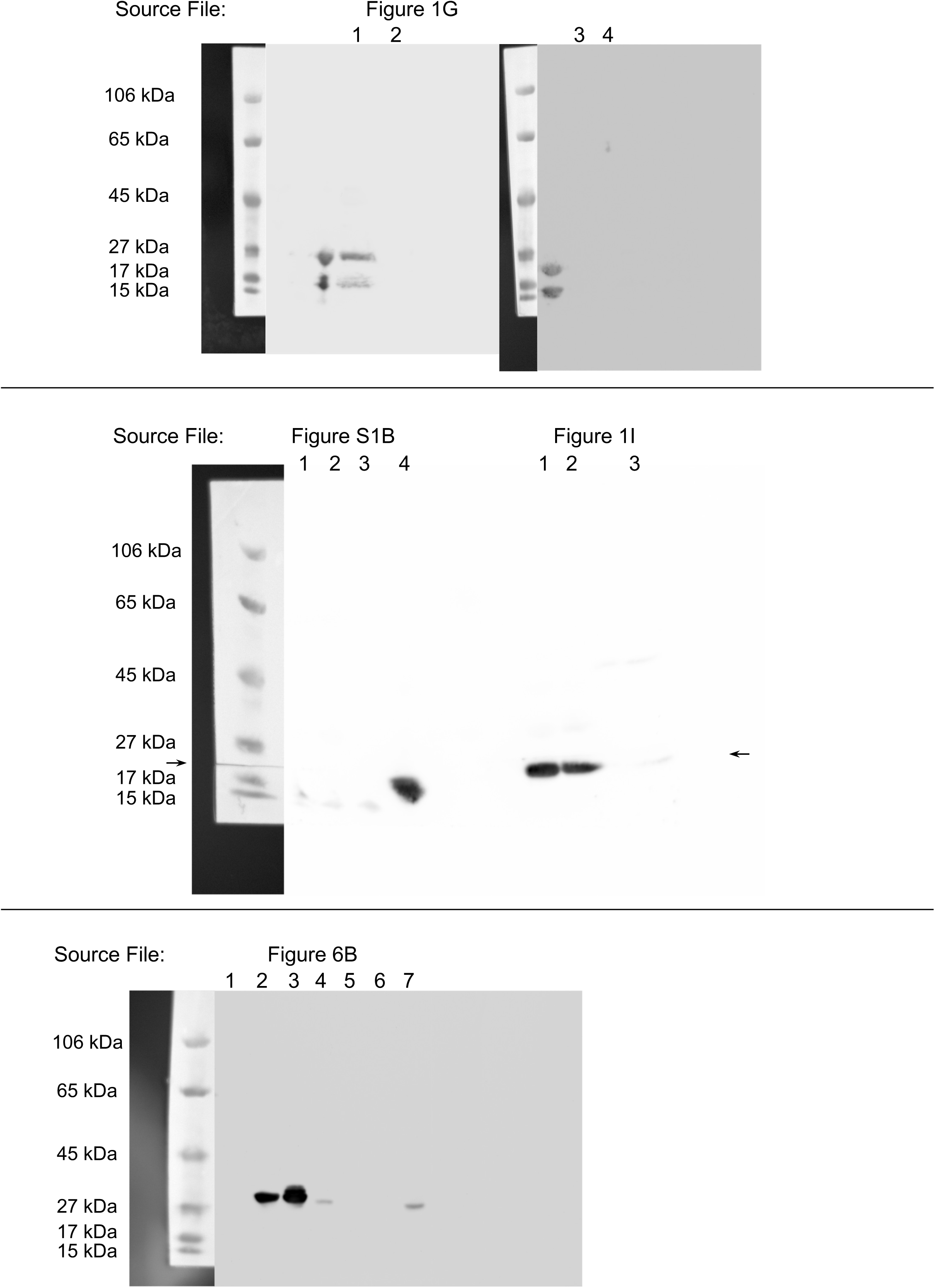

**Table S1.**
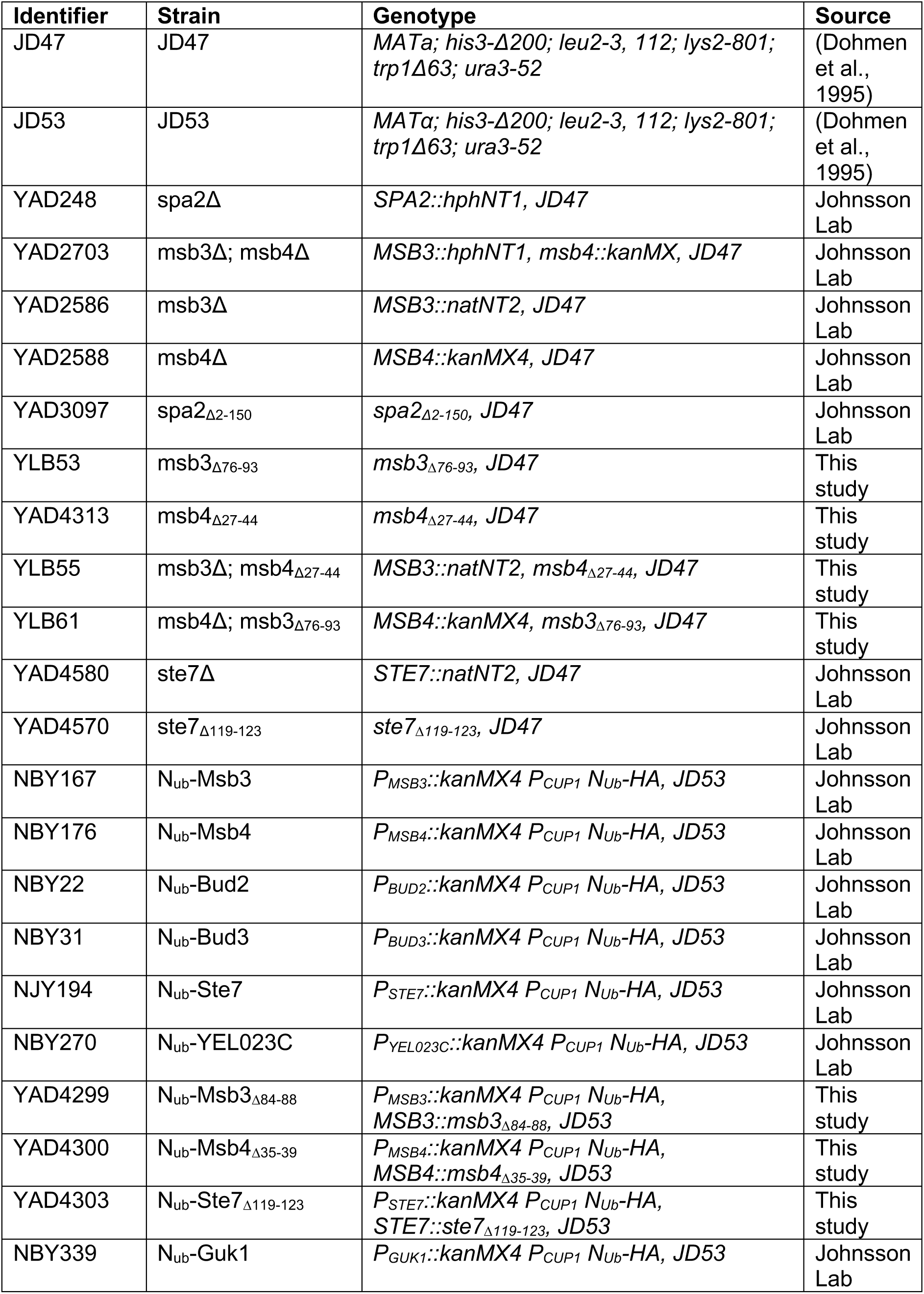
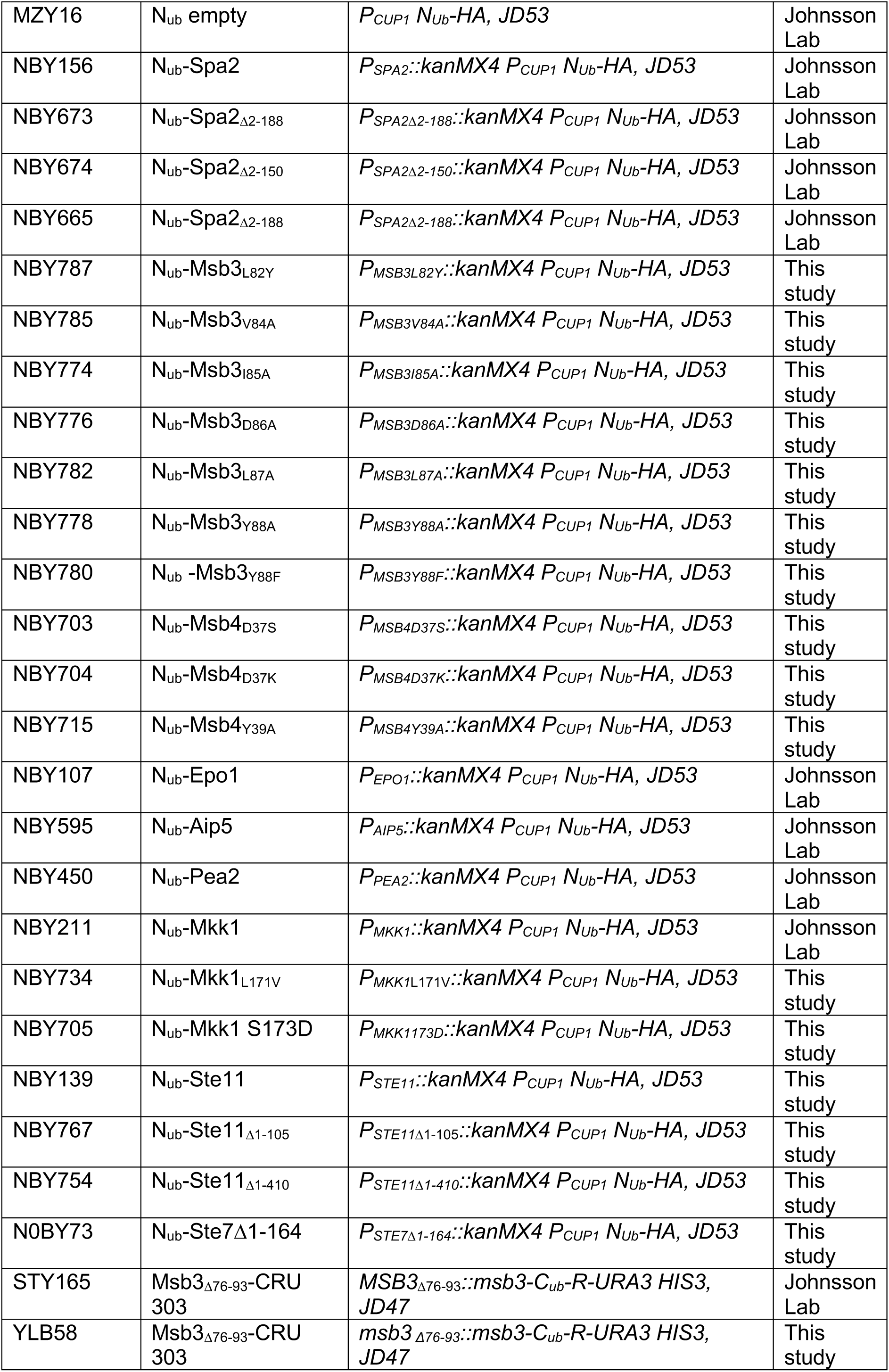
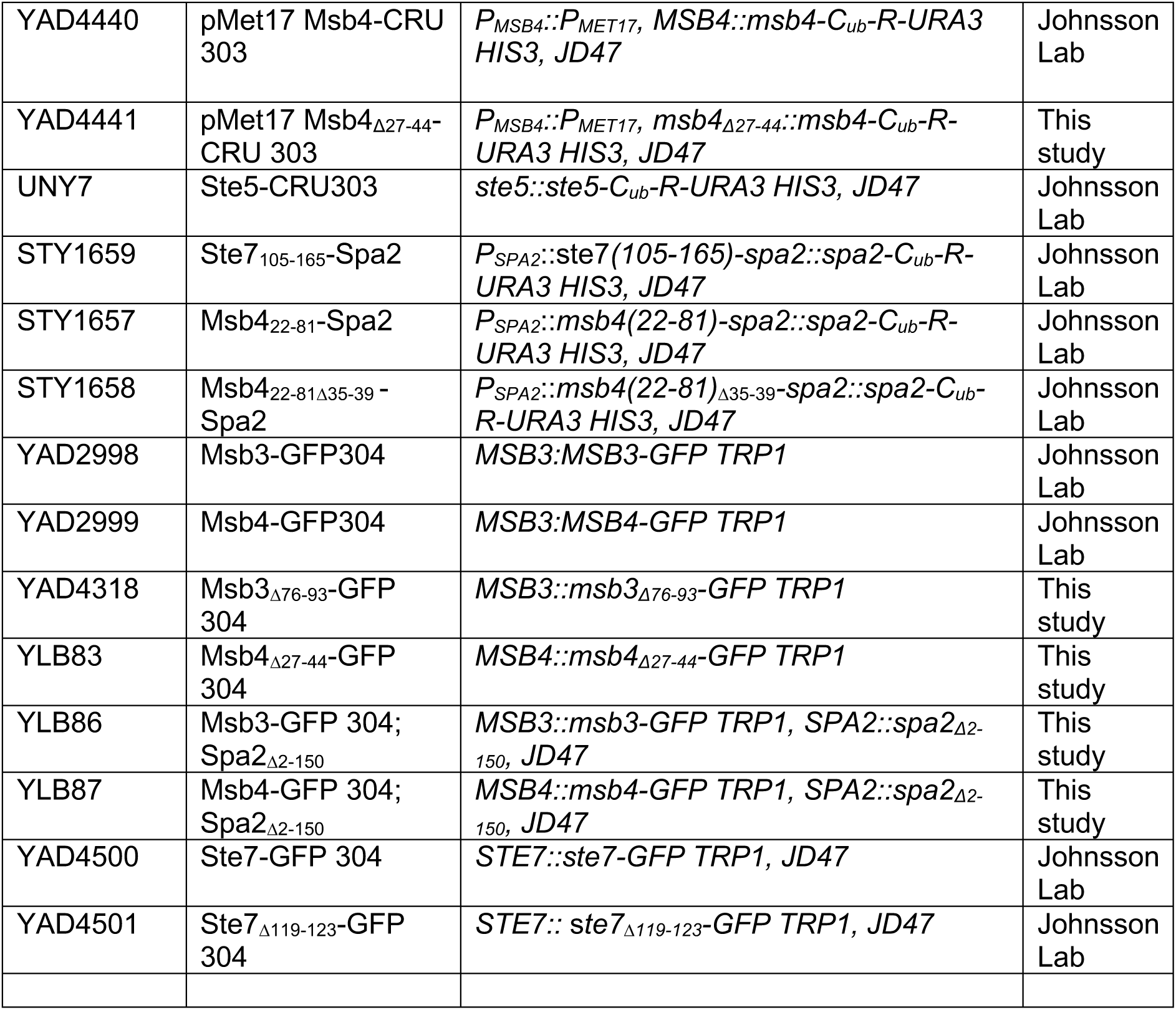
List of *S. cerevisiae* strains used and created in this study.

**Table S2.** List of plasmids used and created in this study.

| Identifier | Plasmid | Genotype | Source |
| --- | --- | --- | --- |
| BS4075 | PMET17-Spa2(1-124)-CRU pRS313 | *PMET17-spa2(1-124)- Cub-R-URA3, AmpR, HIS3, CEN* | Johnsson Lab |
| BA2009 | PMET17-Spa2 L48,51A (1-124) - CRU pRS313 | *PMET17-spa2 L48,51A (1-124)- Cub-R-URA3, AmpR, HIS3, CEN* | Johnsson Lab |
| BL1001 | PMET17-Spa2 L101A (1-124)- CRU pRS313 | *PMET17-spa2 L101A (1-124)- Cub-R-URA3, AmpR, HIS3, CEN* | This study |
| BL1002 | PMET17-Spa2 L109A (1-124)- CRU pRS313 | *PMET17-spa2 L109A (1-124)- Cub-R-URA3, AmpR, HIS3, CEN* | This study |
| BL1003 | PMET17-Spa2 L101,109A (1-124)- CRU pRS313 | *PMET17-spa2 L101,109A (1-124)- Cub-R-URA3, AmpR, HIS3, CEN* | This study |
| BL1004 | PMET17-Spa2 K97D (1-124)- CRU pRS313 | *PMET17-spa2 K97D (1-124)- Cub-R-URA3, AmpR, HIS3, CEN* | This study |
| BA2075 | Msb4(22-81)Δ35-39-Spa(1-124)-CRU | *P<sub>MET17</sub>MSB4(22-81)Δ35-39-SPA2(1-124)-CRU313* | This study |
| BA2080 | Msb4(22-81)Y39A-Spa2(1-124)-CRU | *P<sub>MET17</sub>MSB4(22-81)Y39A-SPA2(1-124)-CRU313* | This study |
| BA2072 | Mkk1(105-164)-Spa2(1-124)-CRU | *P<sub>MET17</sub>MKK1(105-164)-SPA2(1-124)-CRU313* | This study |
| BA2045 | Ste7(105-165)-Spa2(1-124)-CRU | *P<sub>MET17</sub>STE7(105-165)-SPA2(1-124)-CRU313* | This study |
| BA2044 | Msb3(71-130)-Spa2(1-124)-CRU | *P<sub>MET17</sub>MSB3(71-130)-SPA2(1-124)-CRU313* | This study |
| BA2075 | Msb4(22-81)-Spa2(1-124)-CRU | *P<sub>MET17</sub>MSB4(22-81)-SPA2(1-124)-CRU313* | This study |
| BA884 | N<sub>ub</sub>-Spa2<sub>1-150</sub> | *P<sub>CUP</sub>NubHA-Spa2(1-150) kanMX CEN* | Johnsson Lab |
| BA2006 | N<sub>ub</sub>-Msb3<sub>71-100</sub> | *P<sub>Cup</sub>NubHAKanMX Msb3<sub>71-100</sub> CEN* | Johnsson Lab |
| BA2007 | N<sub>ub</sub>-Msb4<sub>21-50</sub> | *P<sub>Cup</sub>NubHAKanMX Msb4<sub>21-50</sub> CEN* | Johnsson Lab |
| BA2008 | N<sub>ub</sub>-Ste7<sub>105-135</sub> | *P<sub>Cup</sub>NuiHAKanMX Ste7<sub>105-135</sub> CEN* | Johnsson Lab |

**Table S3.** Raw fluorescence intensities of GFP fusions for calculating the fluorescence ratios in Figure 4D.

| **Msb3** |  |  | **Msb3<sub>Δ76-93</sub>** |  |  |
| --- | --- | --- | --- | --- | --- |
| Shmoo tip | Cytoplasm | background | Shmoo tip | Cytoplasm | Background |
| 964,583 | 643,375 | 512,833 | 829,5 | 882,417 | 587,083 |
| 801,583 | 614,292 | 525,167 | 758,182 | 725,909 | 578,909 |
| 778,875 | 643,25 | 525,625 | 746,727 | 758 | 576,273 |
| 878,458 | 668,333 | 524,833 | 752,818 | 742,818 | 595,727 |
| 769,5 | 612,625 | 529,375 | 732,818 | 711,909 | 567,636 |
| 973,875 | 689,583 | 528 | 727,545 | 807 | 564,455 |
| 838,917 | 625,917 | 525,75 | 732,636 | 726 | 586,455 |
| 1035,167 | 694,375 | 531,292 | 755,545 | 762 | 541,727 |
| 966,708 | 676,083 | 531,792 | 813,727 | 820,091 | 576,818 |
| 858,708 | 658,833 | 544,333 | 831,545 | 801,727 | 545,273 |

| **Msb4** |  |  | **Msb4<sub>Δ27-44</sub>** |  |  |
| --- | --- | --- | --- | --- | --- |
| Shmoo tip | Cytoplasm | background | Shmoo tip | Cytoplasm | background |
| 1363,364 | 1094,364 | 788,818 | 1294,636 | 1343 | 835,273 |
| 1162,545 | 966,455 | 775 | 1317,273 | 1264 | 824,818 |
| 1442 | 1051,545 | 769,455 | 1112 | 1096,727 | 866,273 |
| 1279,818 | 1089 | 751,909 | 1212,909 | 1306,273 | 776,818 |
| 1435,364 | 1096,091 | 727 | 1212,818 | 1291,636 | 801,636 |
| 1454,273 | 1088,182 | 773,182 | 1257 | 1258,909 | 847,273 |
| 1343,125 | 991,625 | 801,75 | 1268,455 | 1251,727 | 826,818 |
| 1159,438 | 858,455 | 751,545 | 1246,091 | 1397,273 | 834,091 |
| 1642 | 1226,364 | 902,818 | 1209,75 | 1203,182 | 843,545 |
| 1667,455 | 1453,818 | 1069,909 | 1209,545 | 1104 | 856,75 |

| **Ste7** |  |  | **Ste7<sub>Δ112-133</sub>** |  |  |
| --- | --- | --- | --- | --- | --- |
| Shmoo tip | Cytoplasm | background | Shmoo tip | Cytoplasm | background |
| 958,792 | 706,042 | 533,167 | 763,333 | 640,417 | 541,708 |
| 885,75 | 661,375 | 542 | 661,625 | 683,417 | 530,583 |
| 840,083 | 696,917 | 542,581 | 646,708 | 635,042 | 543,208 |
| 909 | 666,25 | 540,292 | 702,167 | 699,667 | 538,5 |
| 785,125 | 649,083 | 553,708 | 748,625 | 663,333 | 524,458 |
| 721,167 | 606,792 | 530,625 | 685,667 | 664,583 | 529,458 |
| 981,208 | 672,833 | 541,458 | 658,083 | 668,5 | 531,042 |
| 926,333 | 627,292 | 545,875 | 687,75 | 638,792 | 537,333 |
| 908,167 | 651,583 | 544,708 | 696,708 | 657,25 | 518,958 |
| 862,125 | 671,875 | 529,833 | 640,125 | 666,083 | 523 |

**Table S4.**
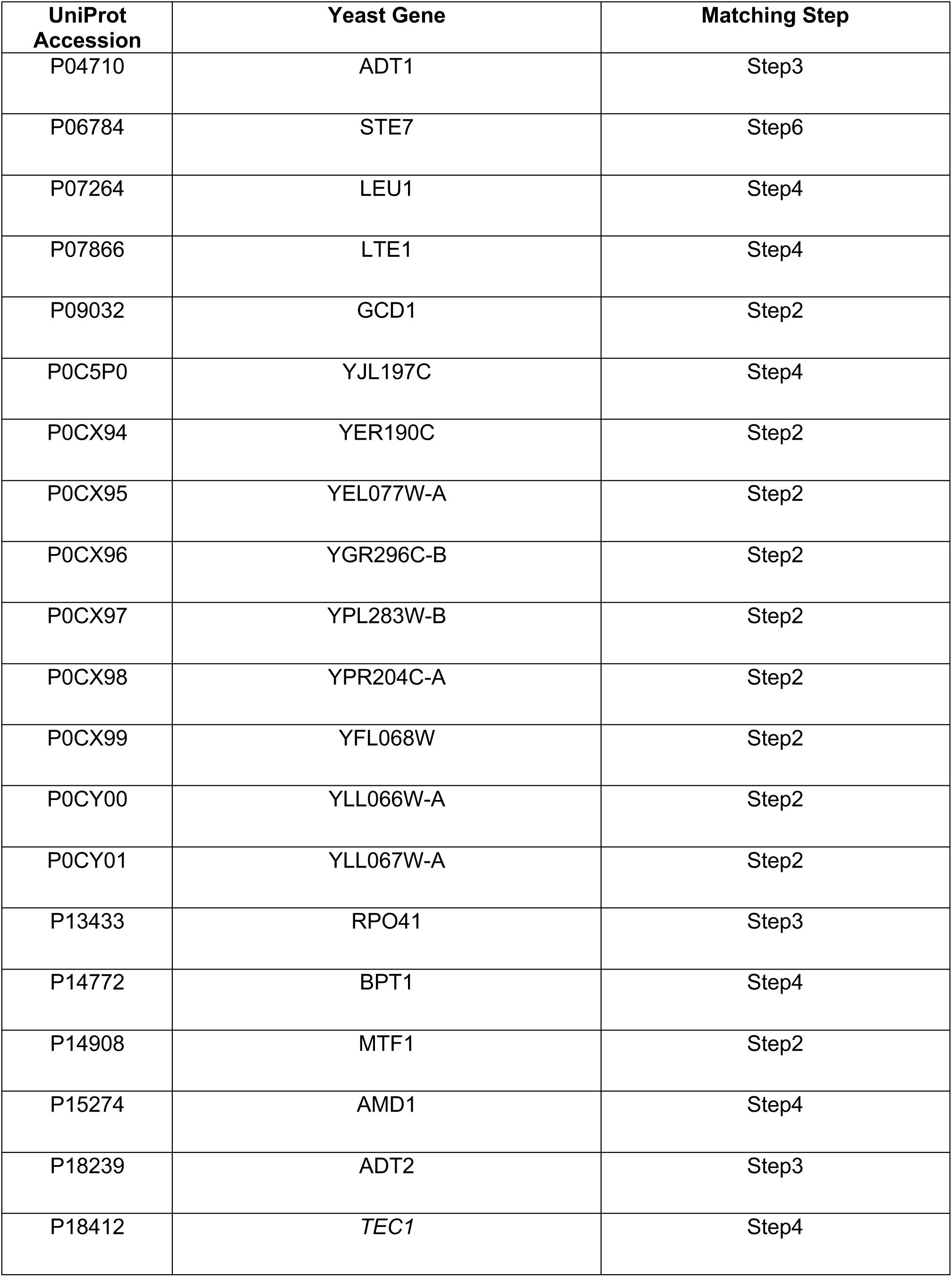
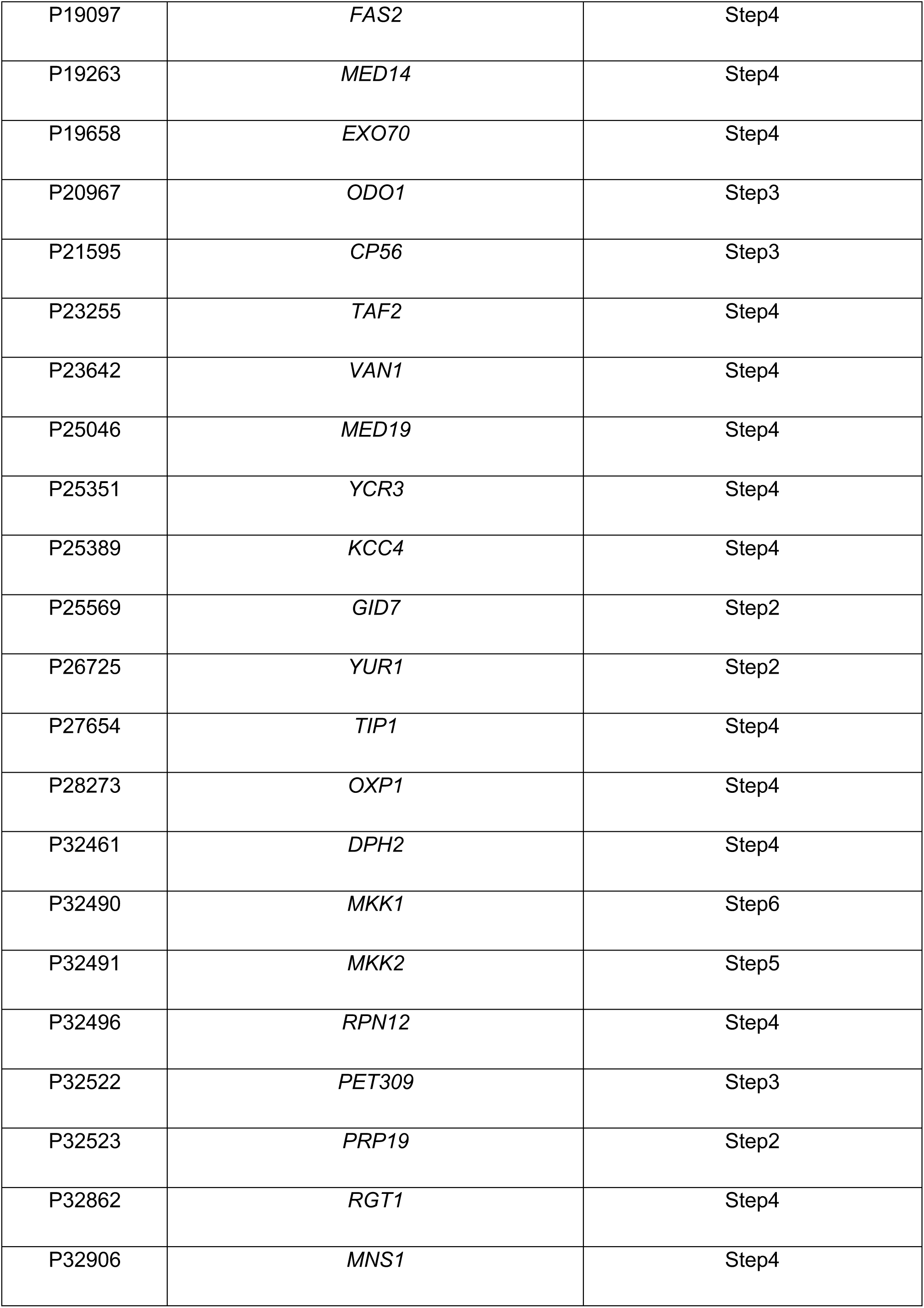
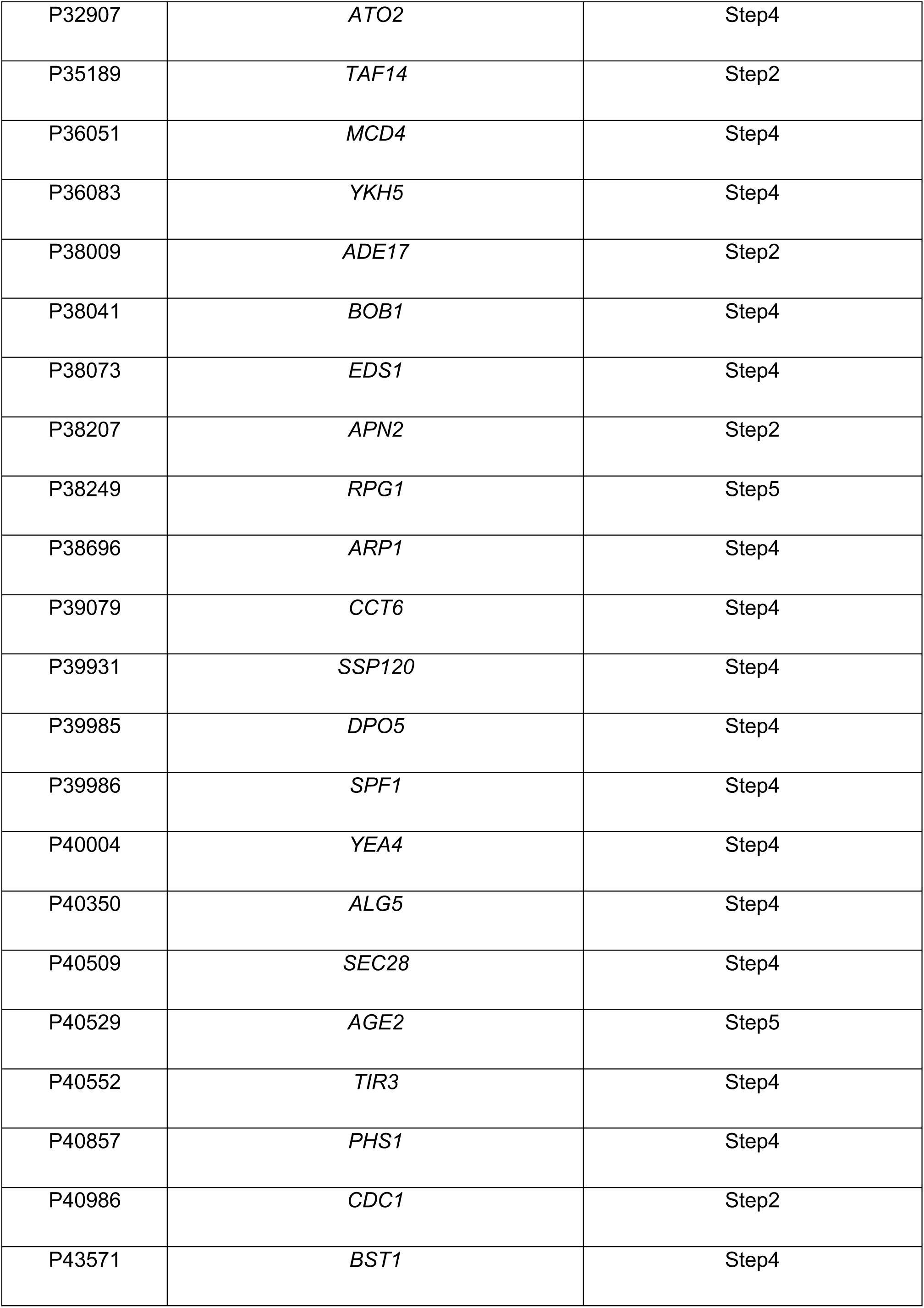
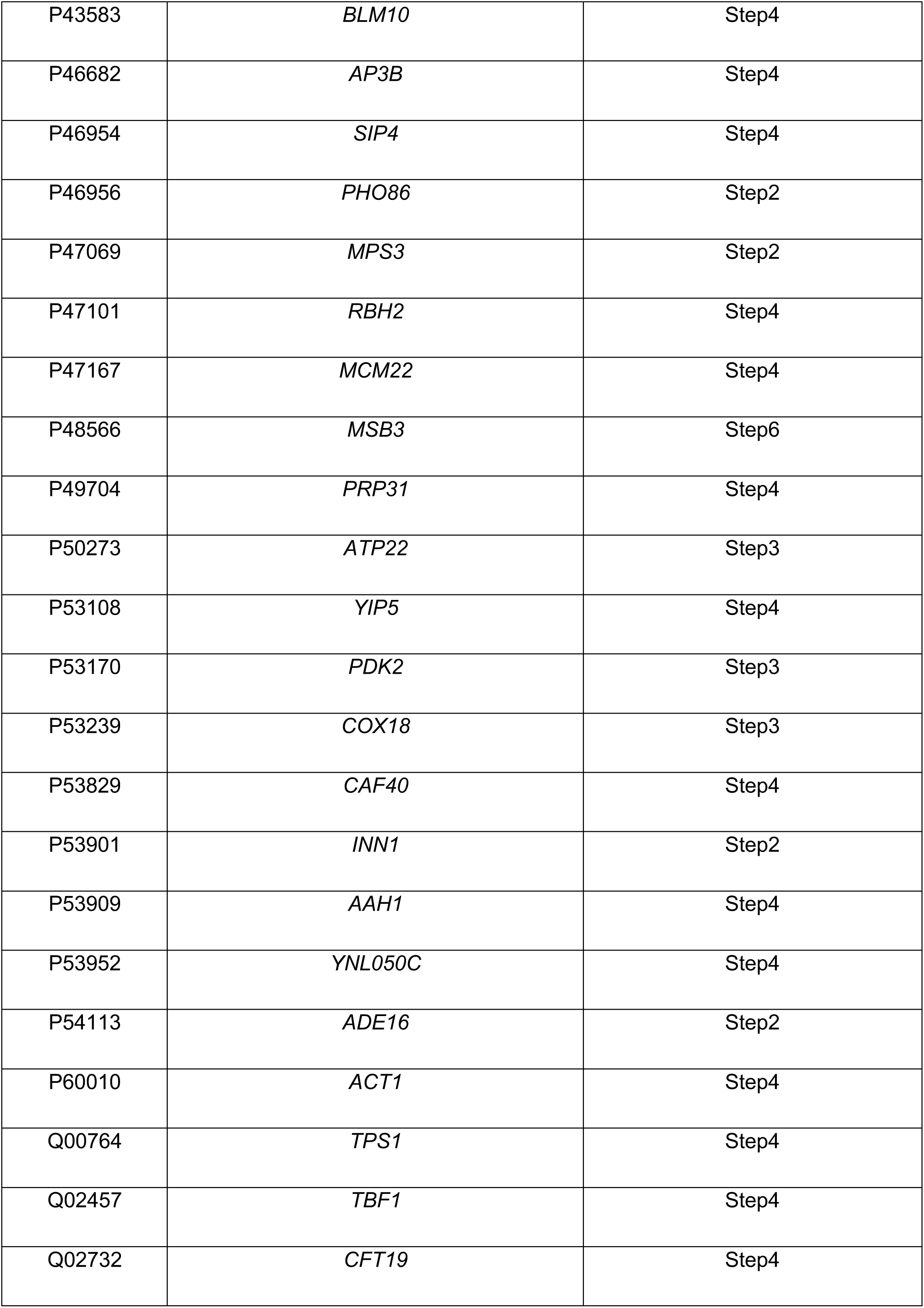
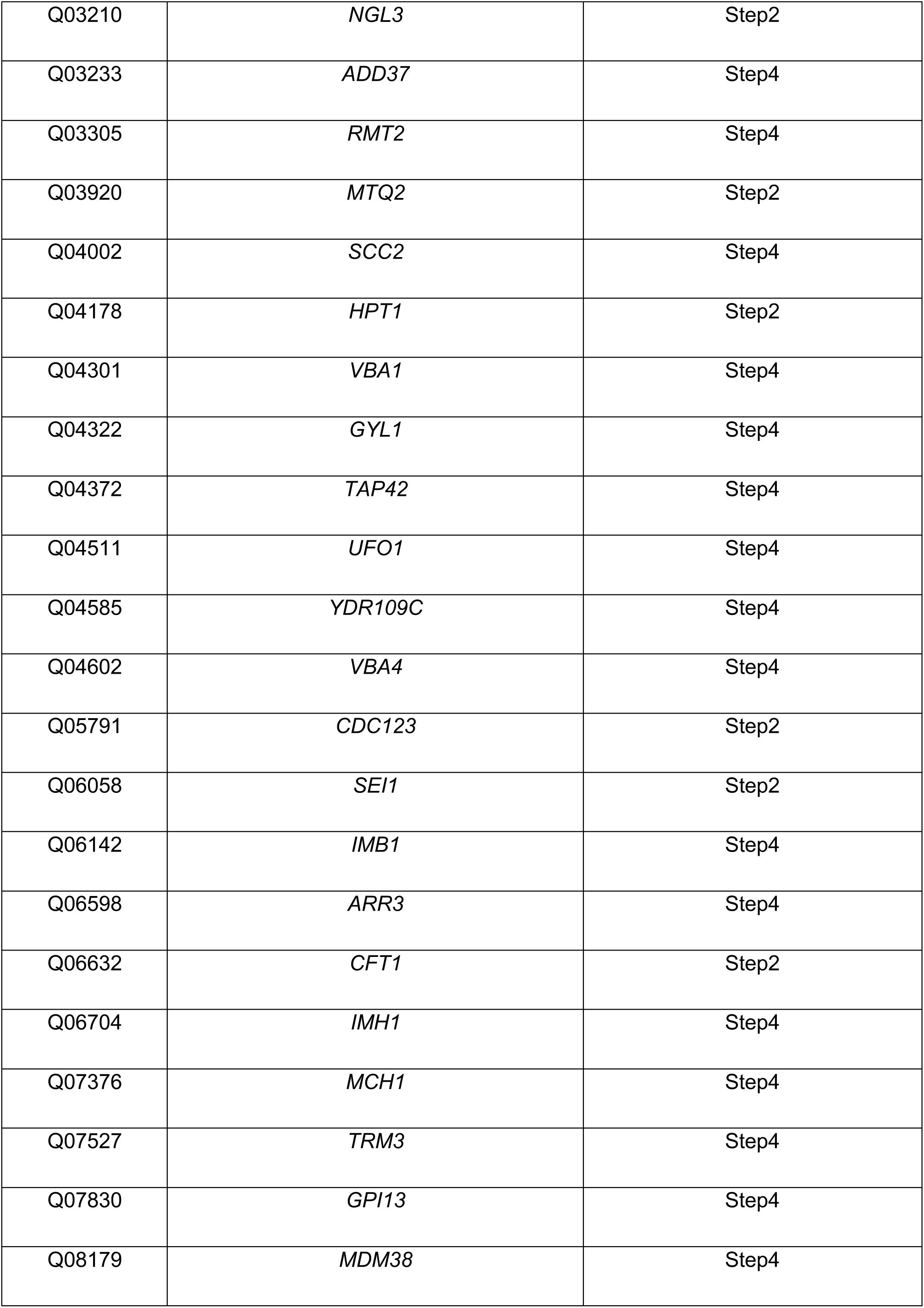
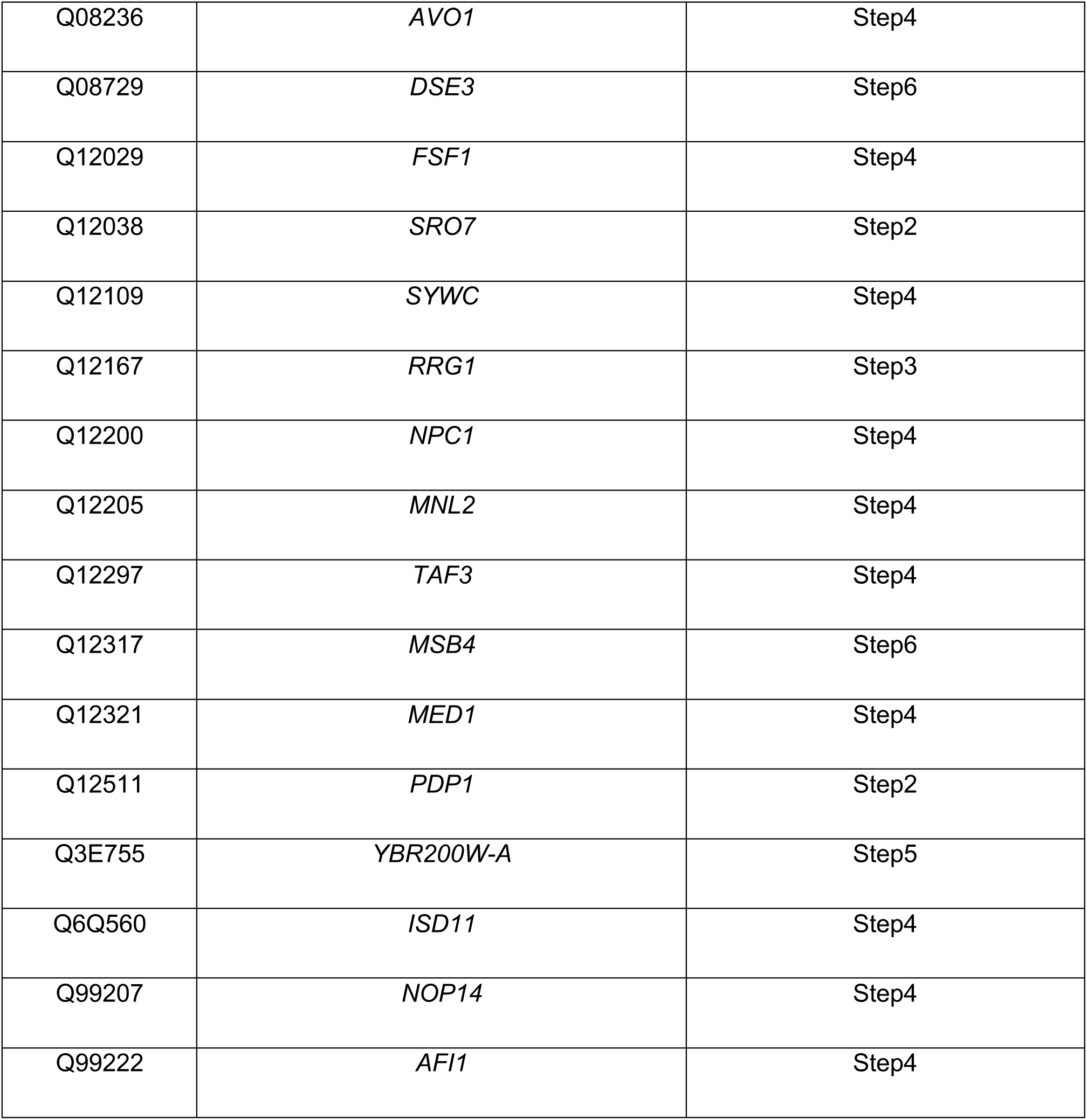
List of potential new FRM-containing binding partners of Spa2.

